# CombinGym: a benchmark platform for machine learning-assisted design of combinatorial protein variants

**DOI:** 10.64898/2026.03.24.714074

**Authors:** Yongcan Chen, Lihao Fu, Xuchao Lu, Wenzhuo Li, Yuan Gao, Yibo Wang, Zhicheng Ruan, Tong Si

## Abstract

Combinatorial mutagenesis is essential for exploring protein sequence-function landscapes in engineering applications. However, while large-scale machine learning benchmarks exist for protein function prediction, they are primarily limited to single-mutant libraries, leaving a critical gap for combinatorial mutagenesis. Here we introduce CombinGym, a benchmarking platform featuring 14 curated combinatorial mutagenesis datasets spanning 9 proteins with diverse functional properties including binding affinity, fluorescence, and enzymatic activities. We evaluated nine machine learning algorithms from five methodological categories (alignment-based, protein language, structure-based, sequence-label, and substitution-based) across multiple prediction tasks, assessing both zero-shot and supervised learning performance using Spearman’s ρ and Normalized Discounted Cumulative Gain metrics. Our analysis reveals the substantial impact of measurement noise and data processing strategies on model performance. By implementing hierarchical dataset splits (0-vs-rest, 1-vs-rest, 2-vs-rest, and 3-vs-rest scenarios), we demonstrate the value of lower-order mutation data for empowering machine learning models to predict higher-order mutant properties. We validated this capacity through both *in silico* simulation (improving fluorescence brightness of an oxygen-independent fluorescent protein) and experimental validation (engineering enzyme substrate specificity), achieving a substantial increase in specific activity. All datasets, benchmarks, and metrics are available through an interactive website (https://www.combingym.org), facilitating collaborative dataset expansion and model development through integration with automated biofoundry platforms.

## Introduction

Engineering proteins with enhanced or novel functions provides new solutions for industrial catalysis, drug development, environmental protection, agricultural engineering, and basic research (1-4). However, predicting how multiple simultaneous mutations affect protein function remains a fundamental challenge due to the nonlinear interactions among amino acid residues that create rugged sequence-function landscapes. These interactions, known as epistatic effects or epistasis, mean that the impact of one mutation often depends on the presence of other mutations (5-7). Epistasis constrains the effectiveness of both rational design and directed evolution strategies for protein engineering (8,9).

Focused combinatorial mutagenesis has been increasingly applied to navigate complex protein sequence-function landscapes (10-16). Recent advances in chip-based oligo synthesis, modular DNA assembly, and high-throughput functional screening now enable creation and profiling of variant libraries containing specific combinations of key residues. This approach concentrates experimental efforts on regions most likely to impact function, guided by structural information, evolutionary conservation, computational predictions, or previous functional screening. However, exploration of the vast combinatorial space remains experimentally intractable, necessitating computational approaches to guide efficient landscape navigation (12,15,17-19). Machine learning (ML) has demonstrated success in modeling protein landscapes by extracting patterns from sequence, structural, and functional data using diverse algorithms (20,21).

As this field produces new protein modeling algorithms at an accelerating pace, standardized benchmarks are essential to evaluate model performance and guide innovation—similar to how ImageNet revolutionized computer vision and how CASP transformed protein structure prediction (22). Several benchmark studies have evaluated ML model performance for predicting protein expression, stability, fluorescence, and protein-ligand interactions (23-29). However, these benchmarks are mainly limited to single-mutant libraries, leaving a critical gap for combinatorial mutagenesis. Furthermore, existing benchmarks rarely include experimental validation, limiting assessment of model overfitting, training data leakage, and extrapolation capabilities (30).

To address these limitations, we present CombinGym (Figure 1), a benchmarking platform specifically designed for function prediction and sequence design of combinatorial protein variants. CombinGym comprises 14 deep mutational scanning (DMS) datasets with over 400,000 characterized variants, spanning diverse functional properties including binding affinity, protein fluorescence, and enzymatic activities. We evaluate nine computational models spanning five methodological categories under both zero-shot and supervised learning scenarios, with special emphasis on predicting higher-order mutant properties from lower-order training data. Our analysis examines critical factors affecting model performance, validates the platform through both computational simulation and experimental testing, and provides all resources through an interactive website (https://www.combingym.org) to facilitate collaborative development.

**Figure 1.**
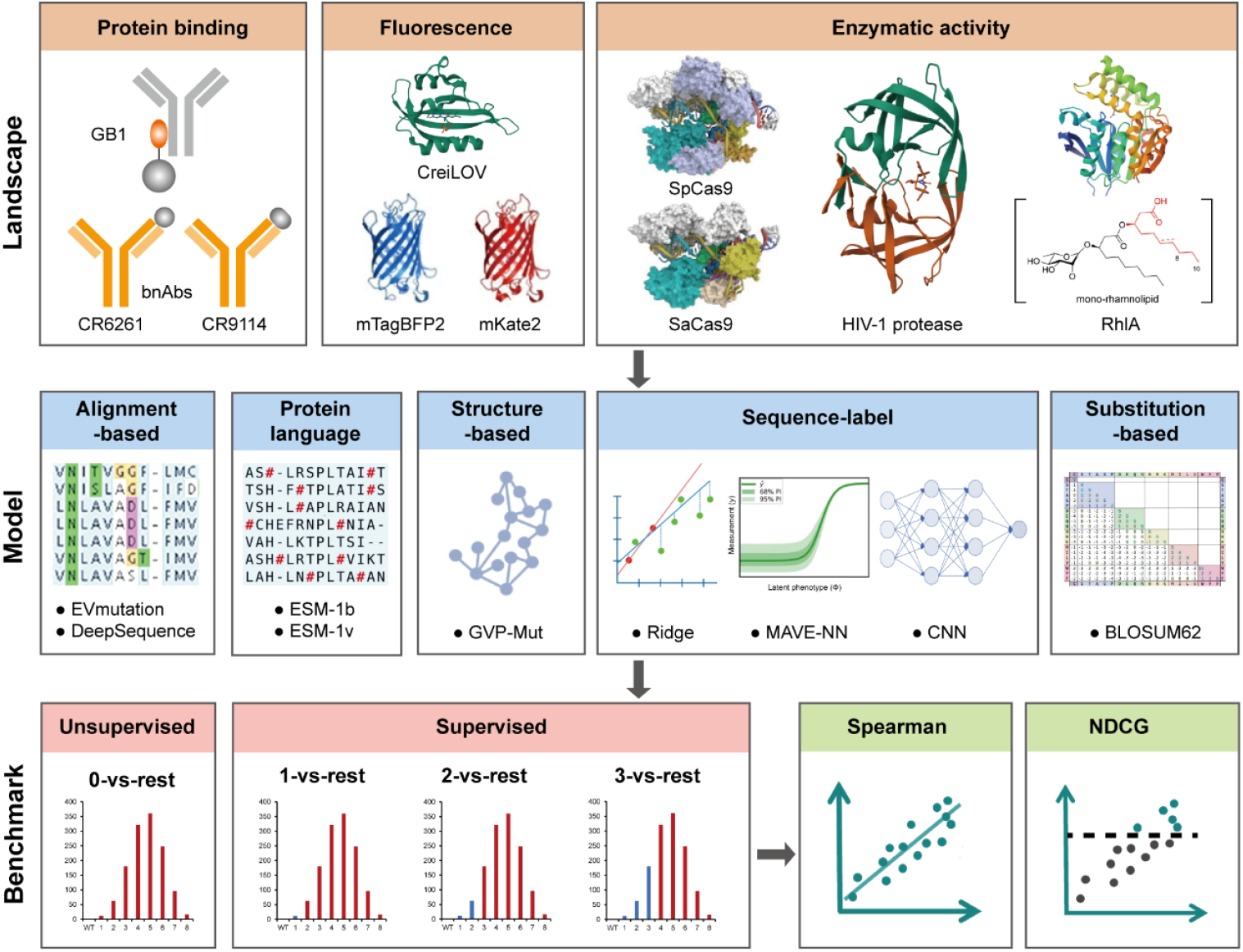
Schematic diagram of the CombinGym benchmarking workflow. CombinGym contains landscapes of nine proteins with the properties spanning from protein binding to fluorescence and enzymatic activity. CombinGym covers baselines ranging from alignment-based, protein language, structured-based, sequence-label to substitution-based models. CombinGym focuses on the benchmarking of protein combinatorial mutants in both unsupervised 0-vs-rest and supervised 1-vs-rest, 2-vs-rest and 3-vs-rest settings, reporting both Spearman and NDCG metrics.

## Materials and Methods

### Deep mutational scanning datasets collection

We compiled and curated 14 published DMS datasets of nine proteins (Table 1). To capture diverse functional contexts, we classified the datasets into three functional categories: protein binding, protein fluorescence, and enzymatic activity.

**Table 1.**
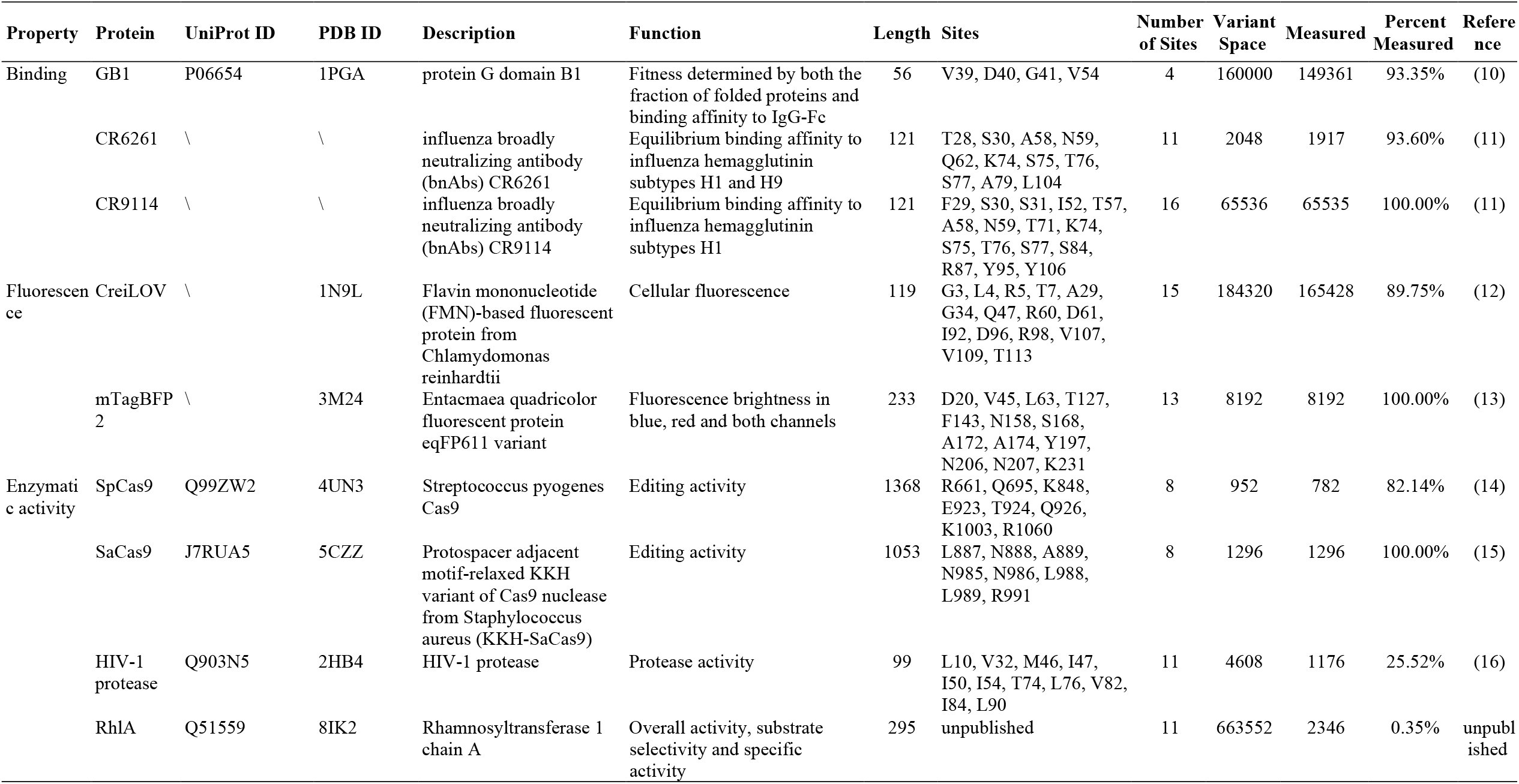
Summary of CombinGym datasets.

### Protein binding

Four landscapes were included for benchmarking on protein binding.

**GB1** is the binding domain B1 of protein G, an immunoglobulin-binding protein in *Streptococcal* bacteria (31,32). The dataset contained 149,361 variants (93.4% of theoretical 20^4^=160,000) across four amino acid sites (V39, D40, G41 and V54) (10). Because over 96% of mutants have fitness value below 0.5, whereas the fitness of wild type is defined as 1 and non-functional variant as 0, we down-sampled non-functional sequences prior training (following the benchmark approach of FLIP (27)) to prevent spurious high-scoring predictions for low-fitness mutants. This treatment resulted in inclusion of all 5,822 sequences with fitness above 0.5 and 2,911 randomly sampled sequences with fitness less than or equal to 0.5.

**CR9114 and CR6261** are two naturally isolated influenza broadly neutralizing antibodies (bnAbs) with 16 or 11 heavy-chain mutations, respectively (33-35). The original studies measured equilibrium dissociation constants of all possible evolutionary intermediates back to the unmutated germline sequences (2^16^=65,536 or 2^11^=2,048) (11). Three (H1, H3 and influenza B) or two (H1 and H9) antigen subtypes were selected to assay equilibrium binding affinities against CR9114 or CR6261, respectively. For CR9114, we utilized only the H1 landscape because H3 and influenza B landscapes were mainly characterized by low fitness values, limiting the information content for model training.

### Fluorescence

Fluorescent protein datasets were selected to represent both oxygen-dependent and oxygen-independent systems with different chromophore chemistries. Four landscapes were included for protein fluorescence benchmarking:

**CreiLOV** is an oxygen-independent, flavin mononucleotide-based fluorescent protein (FbFP) from *Chlamydomonas reinhardtii* (36). We previously assayed the in vivo fluorescence intensities of 165,428 out of 184,320 possible combinatorial mutants among 20 mutations at 15 residues, which were selected based on prior site-saturation mutagenesis screening to identify key residues with functional impact (12).

**mTagBFP2 and mKate2** are the bright blue and bright deep-red variants, respectively, of *Entacmaea quadricolor* fluorescent protein eqFP611 (37-39). They differ by 13 mutations. The published dataset reports blue, red and combined fluorescence of all 2^11^mutants representing the evolutionary intermediates linking these two variants (13), providing a unique benchmark on multiple spectral properties within the same protein framework.

### Enzymatic activities

Enzymatic activity datasets were selected to represent diverse catalytic mechanisms and substrate interactions. Six landscapes were used for the benchmarking:

**SpCas9 and SaCas9** are two CRISPR nucleases widely used for genome engineering (40). For SpCas9, the activities of a combinatorial library with 4×2×17×7=952 variants were assayed based on the loss of red fluorescence through gRNAs targeting the RFP gene sequence (14). For SaCas9, the editing activities of 1,296 combinatorial variants (the product of 12 mutation combinations at WED domain, multiplied by 108 mutation combinations at PI domain) for KKH-SaCas9 were measured using a GFP reporter system (15). These nucleases were selected because their complex multi-domain structure and multistep catalytic mechanism present substantial modeling challenges. **HIV-1 protease** is essential for viral maturation and is an important target of combination antiretroviral therapy (41). The dataset quantified the relative enzymatic activity of 2,736 out of 4,608 (2^9^×3^2^) possible combinatorial variants covering 11 resistance-associated sites (16).

**RhlA** catalyzes the formation of the lipid moiety of rhamnolipids (RLs) (42). Our group has previously constructed a subset of combinatorial RhlA library consisting of 30 selected amino acid substitutions at 11 residues with a theoretical diversity of 663,552 (unpublished). Measured phenotypes include the overall activity, specificity, and specific activity of variants with 1 to 3 mutations.

### DMS dataset normalization

To enable fair comparison across datasets with different measurement scales and distributions, we performed min-max normalization to transform all measurement values into a range between 0 and 1 to ensure the scale uniformity across datasets. The normalization formula is:

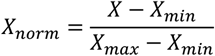

Where *X* represents the original measurement values, *X*_*min*_ is the minimum value in the dataset and *X*_*max*_ is the maximum value. This approach preserves the relative ranking of variants while standardizing the numerical range across datasets.

### Data Splitting for higher-order prediction

The central goal of CombinGym is to evaluate model performance in predicting higher-order mutant properties from lower-order training data. We implemented a hierarchical splitting strategy to progressively increase the mutational complexity in test sets while training on simpler variants:

#### 0-vs-rest

Zero-shot prediction scenario, where models must predict all mutant phenotypes without any training on the measured data. This evaluates baseline predictive capability using EVmutation, DeepSequence, ESM-1b, ESM-1v and BLOSUM62 models.

#### 1-vs-rest

Models are trained on wild-type and single mutants only, then evaluated on their ability to predict double, triple, and higher-order mutant properties. This tests the ability to extrapolate from minimal training data.

#### 2-vs-rest

Models are trained on WT, single, and double mutants, then evaluated on triple and higher-order variants. This represents a common practical scenario in protein engineering where combinatorial libraries are designed based on screening of single and double mutants.

#### 3-vs-rest

Models are trained on WT through triple mutants, then evaluated on higher-order (>3) combinations. This tests whether including triple mutant data greatly improves prediction of more complex combinations. For RhlA datasets, 3-vs-rest settings are absent since the available data only includes variants with up to three mutations.

### Multiple sequence alignment (MSA)

For MSA-based models (EVmutation and DeepSequence), alignments were generated using the EVcouplings framework (43) on a local server. For each target protein, the WT sequence (Table 1) was used as the query, and the alignments were searched against the UniRef100 dataset. We employed the default parameters on the EVcouplings webserver (search iterations = 5, position filter = 70%, sequence fragment filter = 50%, removing similar sequences = 100%, downweighting similar sequences = 80%) with the exception of bitscore thresholds, which were varied to assess MSA quality impact. Initial bitscore thresholds were set to 0.1, 0.3, 0.5, and 0.7, and then further optimized based on two criteria: (1) the redundancy-reduced number of sequences and (2) the coverage of targeted combinatorial sites. This MSA generation approach allowed us to investigate how MSA quality and depth affect model performance—an important consideration given the variable evolutionary conservation of different proteins in our benchmark.

### Protein structure prediction

Because experimentally solved structures are unavailable or incomplete for certain proteins, we employed AlphaFold3 (44) to predict the 3D structures of the target sequence of all DMS assays, instead of retrieving PDB files from the RCSB Protein Data Bank (45). The incorporation of AlphaFold3 also standardized the generation of protein structures and presented an end-to-end workflow directly from sequence to function. The wild-type sequences used for DMS were taken as the input.

### Baseline models

We selected nine baseline models representing diverse approaches to protein fitness prediction, categorized into five main methodological groups.

### Alignment-based models

**EVmutation** (46) predicts mutation effects by capturing pairwise residue dependencies between positions using a Potts model derived from MSA. This statistical approach explicitly models co-evolutionary patterns among amino acids within protein sequences and has been widely applied to predict the effects of single mutations.

**DeepSequence** (47) extends beyond pairwise dependencies by implementing a deep generative model based on a variational autoencoder (VAE) framework. Compared to EVmutation, DeepSequence can capture higher-order, non-linear dependencies in biological sequences through its latent variable representation.

### Protein language models

**ESM-1b** (48) and **ESM-1v** (49) are both transformer-based models with 750 M parameters, trained with the masked language modeling objective on UniRef50 and UniRef90 datasets, respectively. For supervised prediction tasks, we implemented these models within the FLIP framework using a batch size of 256, a learning rate of 0.001, and the Adam optimizer (27). We employed the per-amino-acid embedding pooling method, which enables the supervised model to utilize a 1D attention layer for dynamic sequence pooling to enhance regression prediction.

### Structure-based model

**Geometric vector perceptron (GVP)-Mut** model (50) was included as the sole structure-based method in our benchmark. This approach utilizes a 3D graph neural network to extract features from protein structures. The architecture builds on GVP-GNN (51), incorporating only the backbone in the graph neural network, as the sidechains were not precisely predicted by AlphaFold3.

### Sequence-label models

**Convolutional neural network (CNN)** and **Ridge** regression were implemented following the FLIP framework architecture (27), using one-hot encoding of amino acid sequences. The CNN architecture consists of a convolution layer with a kernel width of 5 and 1024 channels, a ReLU nonlinearity, a 2048-dimensional linear mapping, a maximum pooling over the sequence, and a 1-dimensional linear mapping. For ridge regression, we employed the scikit-learn implementation with default parameters.

**MAVE-NN** (52) implements a neural-network-based framework that uses an information-theoretic approach for learning genotype-phenotype maps from multiplex assays of variant effect datasets. A key feature of MAVE-NN is its explicit modeling of the latent phenotype specified by the genotype-phenotype map, which serves as input to a measurement process that includes a global epistasis (GE) component and a noise model. Based on our previous research (12), we utilized the blackbox genotype-phenotype map model coupled with GE regression and skewed-*t* noise models, trained with a learning rate of 0.001, 500 epochs, batch size of 64, and early stopping patience of 25.

### Substitution-based model

**BLOSUM62** (53) represents a simple but effective baseline based on evolutionary substitution frequencies. For each mutant sequence, we calculated the total BLOSUM62 substitution score by summing the individual scores for each amino acid substitution between the wild-type and mutant sequences (54). This approach was implemented within the FLIP framework (27) and serves as a reference point for assessing the value of more complex modeling approaches.

### Evaluation metrics

We employed two complementary metrics—Spearman’s correlation coefficient (ρ) and Normalized Discounted Cumulative Gain (NDCG)— to evaluate model performance across different prediction scenarios and applications.

#### Spearman’s correlation coefficient

Spearman’s correlation coefficient (ρ) evaluates how well the ranking of predictions correlates with the ranking of measurements, making it suitable for assessing overall predictive performance across the entire fitness landscape. This non-parametric measure is particularly appropriate for protein fitness data, as it does not assume linear relationships or normal distributions. The formula is:

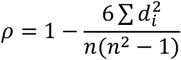

Where *n* is the number of data points and *d*_*i*_ is the difference between the ranks of each pair of corresponding values between prediction and measurement.

#### Normalized Discounted Cumulative Gain (NDCG)

While Spearman’s ρ assesses overall ranking correlation, protein engineering typically focuses on identifying top-performing variants. Therefore, we also calculated the Normalized Discounted Cumulative Gain (NDCG), which specifically measures how well the model identifies high-fitness variants, with stronger weighting for correctly ranking the top-performing mutants. This metric is particularly relevant for protein design applications where researchers aim to prioritize a small subset of variants for experimental testing. The NDCG calculation (55) is:

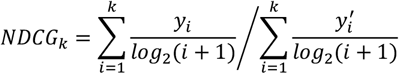

Where *k* defines the rank position up to which NDCG is calculated, *i* denotes the position in the ranking list, *y*_*i*_ means the normalized measurement value of the mutant at position *i* in the predicted ranking and 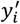 is the normalized measurement value of the mutant at position *i* in the ideal ranking. The discount factor *log*_2_(*i* + 1) reduces the impact of lower-ranked items. In this study, NDCG was calculated for the top 1% of the ranked mutants to focus on the most relevant portion of the prediction for practical applications.

## Data and code availability

All data, code, and resources have been made publicly available to maximize reproducibility and facilitate community adoption:

### Source code

The implementation code for each baseline model, along with our data processing, prediction, and evaluation pipelines, is available in our GitHub repository (https://github.com/sitonglab/CombinGym).

### Dataset access

All original raw and processed DMS datasets are provided in a consistent, standardized format to enable direct comparison across proteins and properties.

### Supporting files

We provide all necessary files for model training and evaluation, including wild-type amino acid sequences (FASTA format), multiple sequence alignments (a2m format), and AlphaFold3-predicted protein structures (PDB format).

### Results and leaderboard

Detailed performance metrics for each baseline model, landscape, and benchmark scenario are available through our interactive website (https://www.combingym.org/), which also includes a continuously updated leaderboard for community contributions.

These resources establish CombinGym as a foundation for collaborative development and rigorous evaluation of machine learning approaches for combinatorial protein engineering.

## Results

### Datasets and benchmark strategies overview

CombinGym comprises nine proteins spanning diverse functional categories: enzymes (SaCas9 and SpCas9 endonuclease, HIV-1 protease, RhlA acyltransferase), fluorescent proteins (eqFP611 and CreiLOV), and protein-binding proteins (bnAbs CR6261 and CR9114, and GB1) (Figure 1; Table 1). Two datasets contain multiple functional readouts: the RhlA dataset contains multiple phenotypes including enzyme activity and substrate selectivity, while the eqFP611 dataset (13) encompasses blue, red and combined fluorescence measurements.

We analyzed the distribution of mutants by mutation number and their corresponding fitness distributions. Mutant count distribution exhibited normal or near-normal distributions, except for RhlA and GB1 (Supplementary Figure S1). The GB1 dataset contains all 20^4^combinations at four selected sites, while the RhlA dataset covers only single, double, and triple mutants within a sequence space of 30 mutations at 11 sites. The overall fitness distribution generally decreases with increasing mutation number for most landscapes (Supplementary Figure S2). However, distinct patterns emerged: with increasing mutation number, the specificity and specific activity phenotypes of RhlA show initial increases followed by decreases, the red fluorescence of eqFP611 gradually increases, the combined fluorescence first decreases then increases, and the bnAbs binding affinities all show continuing increases. These diverse landscape topologies provide varied benchmarks for model evaluation.

Our benchmarking framework (Figure 1) evaluates nine baseline models spanning five methodological categories: MSA-based models (EVmutation (46) and DeepSequence (47)), protein language models (ESM-1b (48) and ESM-1v (49)), structure-based models (GVP-mut (50)), sequence-label models (MAVE-NN (52), CNN and Ridge), and substitution-based models (BLOSUM62 (27)). EVmutation, DeepSequence and BLOSUM62 are unsupervised methods based on evolutionary information, while ESM-1b and ESM-1v can be used in both unsupervised and supervised learning settings. The remaining models are supervised learning approaches. We evaluated the generalization capability for higher-order mutants through four scenarios: unsupervised 0-vs-rest, and supervised 1-vs-rest, 2-vs-rest, and 3-vs-rest. For each task, we calculated Spearman’s ρ and NDCG metrics to assess performance in protein prediction and design applications.

### Influence of measurement noise on supervised model performance

We first examine how measurement noise affects model performance. Biological replicates were available for all datasets except for RhlA. Pearson’s correlation coefficients between biological replicates revealed distinct patterns: SaCas9 and SpCas9 showed low correlations (Pearson’s r < 0.5), while other proteins exhibited high correlations (Pearson’s r > 0.8) (Figure 2A; Supplementary Figure S3). The low reproducibility of Cas9 proteins is likely due to the complexity of their catalytic mechanism, particularly the use of different sgRNAs between biological replicates. Measurement noise in experimental data degrades prediction accuracy and generalization of machine learning models through multiple mechanisms (56).

**Figure 2.**
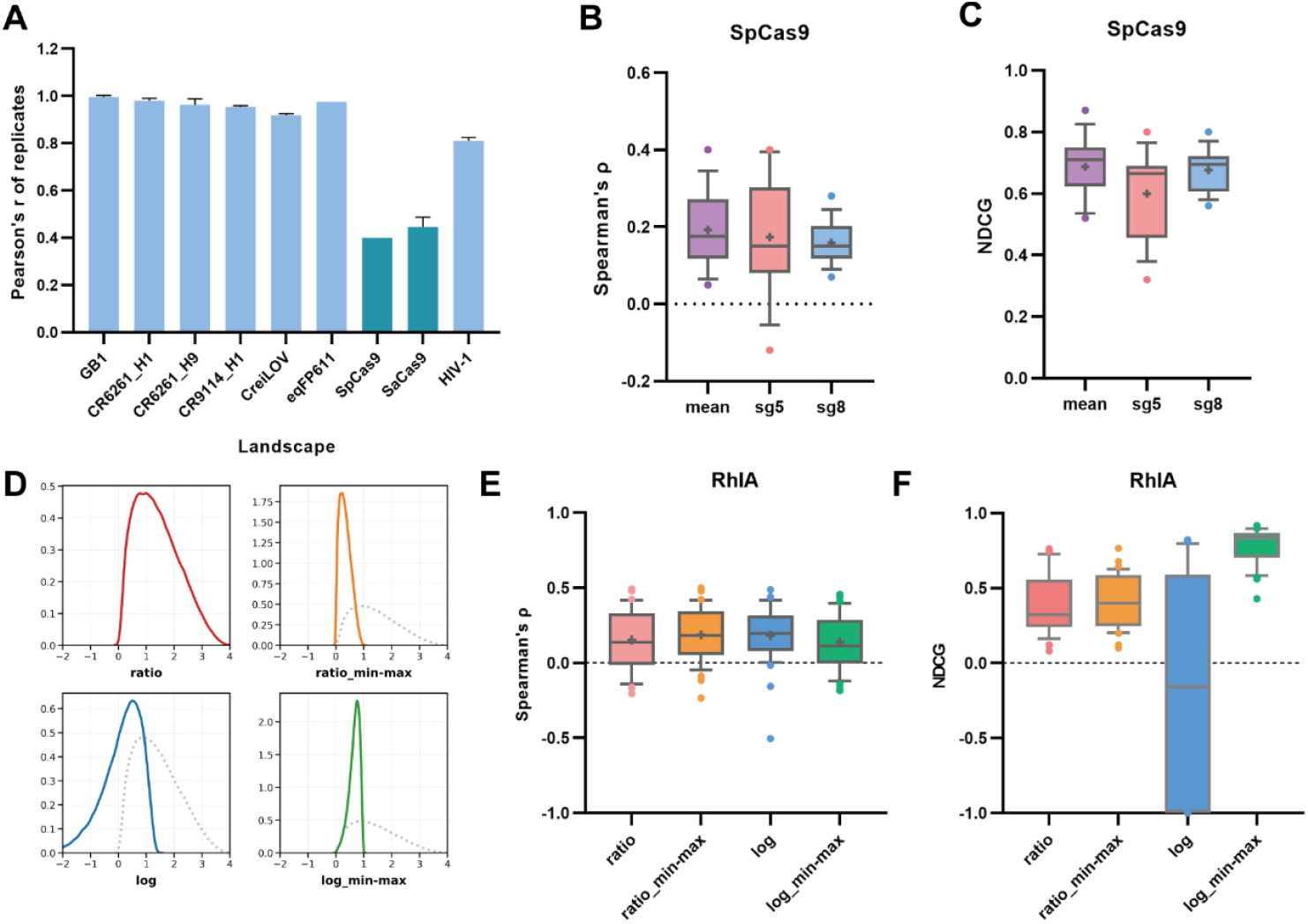
Impact of measurement noise and normalization on supervised model performance. (**A**) The Pearson’s correlation coefficients of the biological replicates for each deep mutational scanning datasets. (**B** and **C**) The Spearman’s ρ (**B**) and NDCG (**C**) of the supervised models trained on the sg5, sg8 or the mean values in SpCas9 datasets. (**D**) A simulation diagram showing the change in data distribution after log, min-max normalization, or both. The dashed lines in ratio_min-max, log or log_min-max indicates the original ratio distribution. (**E** and **F**) The Spearman’s ρ (**E**) and NDCG (**F**) of the supervised models trained on the original ratio or after log, min-max normalization, or both, for RhlA datasets.

The on-target activities on RFPsg5 and RFPsg8 for SpCas9 were measured separately by Thean et al. (15). We trained ESM-1b, ESM-1v, Ridge, CNN, and MAVE-NN models with RFPsg5, RFPsg8 and mean activity as fitness scores separately and compared model performance. Training with sg5 and sg8 datasets yielded distinct model performances, while using the mean values produced better results (Figure 2B and C). ESM-1b, Ridge and MAVE-NN showed greater sensitivity to measurement noise than others, as evidenced by larger performance differences between models trained on mean values versus individual biological replicates (Supplementary Figure S4).

### Influence of normalization on supervised model performance

Functional value calculations vary substantially across datasets due to differences in functional assays and data processing methods. Measurement assays include fluorescence-activated cell sorting, growth selection, affinity-based enrichment, and mass spectrometry. Fitness values were derived through enrichment scores, weighted fluorescence averages across different bins, or nonlinear regression of readouts at various concentrations. Common normalization approaches include WT normalization, log transformation, min-max normalization, and Z-score standardization.

To investigate how data normalization affects machine learning model performance, we used RhlA as an example, where phenotypes were defined as the ratio of mutants to wild-type. We applied log transformation, min-max normalization, or both (Figure 2D), and trained ESM-1b, ESM-1v, Ridge, CNN and MAVE-NN models for 0-vs-rest, 1-vs-rest, and 2-vs-rest tasks. Overall, min-max normalization resulted in higher Spearman’s ρ, and combining it with log normalization increased the NDCG metrics (Figure 2E and F; Supplementary Figure S5). Log transformation alone can lead to negative NDCG, indicating that min-max normalization is essential, particularly when negative fitness values are present after log normalization. Different models exhibit varying sensitivities to min-max normalization (Supplementary Figure S6), suggesting that min-max normalization is beneficial for comparing datasets with distinct scales and models with different algorithms.

### Influence of MSA depth on MSA-based model performance

Machine learning methods based on co-evolution of natural sequences, such as EVmutation(46) or DeepSequence(47), require high-quality multiple sequence alignment data. Alignments yielding >80% coverage of the target domain length and redundancy-reduced sequence counts ≥ 10L (where L is the sequence length) are recommended (46,47). Previous studies showed that DeepSequence is more susceptible to failure than EVmutation when trained on low-quality MSAs (47,55). To evaluate the effects of MSA quality on performance, we generated alignment files with gradient bitscore thresholds for each protein using local EVcouplings (43) (see Methods). MSA depths for SpCas9, SaCas9 and mTagBFP2 proteins were lower than 10 sequences/L when using bitscore thresholds 0.3, 0.5 and 0.7, while the MSA depth of HIV-1 protease was high but insensitive to different bitscore thresholds (Supplementary Figure S7A). A very high MSA depth was achieved for RhlA when using bitscore threshold 0.3, but it decreased sharply with increasing thresholds and became insensitive to further threshold increase. By employing finer-grained bitscore thresholds, we explored and obtained extended MSA depth coverage (Supplementary Table S1).

We selected CreiLOV, bnAbs CR9114 and CR6261, and GB1 proteins to evaluate the impact of MSA depth on the performance of EVmutation and DeepSequence. For each protein, we applied three distinct bitscore thresholds to vary MSA depth. For CreiLOV and bnAbs proteins, MSA depth exceeded the recommended minimum 10 sequences/L (Figure 3A). We used common mutant sets for different bitscore thresholds when varying numbers of mutant sites were covered by MSA. Performance varied considerably across different targets, with Spearman’s correlation coefficients between prediction and measurement being positive for CreiLOV, negative for bnAbs proteins, and near zero for GB1 (Figure 3B-D). However, no differences in model performance emerged across different bitscore thresholds, with Pearson’s r exceeding 0.97 for CreiLOV and bnAbs proteins, and exceeding 0.7 for GB1 (Supplementary Figure S7B and C). DeepSequence performed worse than EVmutation on GB1, supporting previous findings that DeepSequence is susceptible to failure when training on low-quality MSA (47,55). These results indicated that MSA depth does not play a central role in MSA-based machine learning algorithms beyond the minimum threshold.

**Figure 3.**
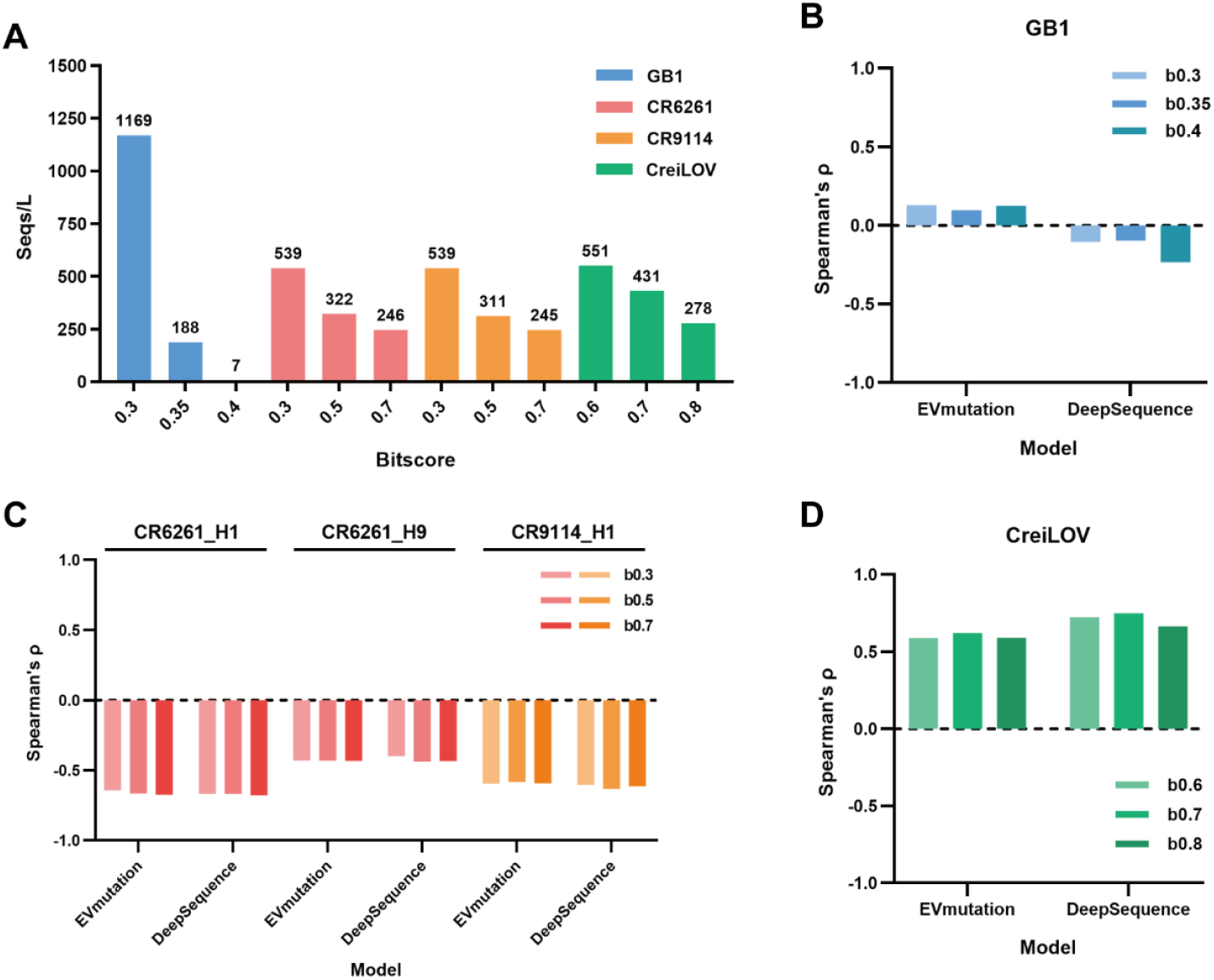
Impact of MSA depth on unsupervised alignment-based models. (**A**) MSA depth and corresponding bitscore thresholds used for GB1, CR6261, CR9114 and CreiLOV. (**B**-**D**) The Spearman’s ρ of EVmutation and DeepSequence for GB1 (**B**), CR6261 and CR9114 (**C**) and CreiLOV (**D**) with gradient MSA depths.

### Model benchmarking

Based on these findings, we performed model benchmarking for higher-order mutant prediction and design. The default bitscore threshold of 0.5 was used except for SaCas9-KKH and CreiLOV (0.7) and for GB1 (0.4). Mutated sites covered by MSA are listed in Supplementary Table S1. We applied min-max normalization to all datasets to ensure consistent scaling. For measurements with biological replicates, we used mean values as fitness scores. We predicted protein structures using AlphaFold3 as inputs for the GVP-Mut model, even when experimentally resolved structures were available, to avoid biases from mutations, deletions, or insertions. We focused on evaluating generalization capability for higher-order mutants through four scenarios: unsupervised 0-vs-rest and supervised 1-vs-rest, 2-vs-rest, 3-vs-rest scenarios. For the RhlA dataset, which contains mutations up to three, we excluded the 3-vs-rest task. We calculated Spearman’s ρ and NDCG metrics for each task to assess performance in protein prediction and design applications. Overall, MAVE-NN and GVP-Mut models achieved top performance for protein prediction across all task scenarios, while GVP-Mut, MAVE-NN, and Ridge regression excelled at protein design (Table 2, Figure 4A).

**Table 2.**
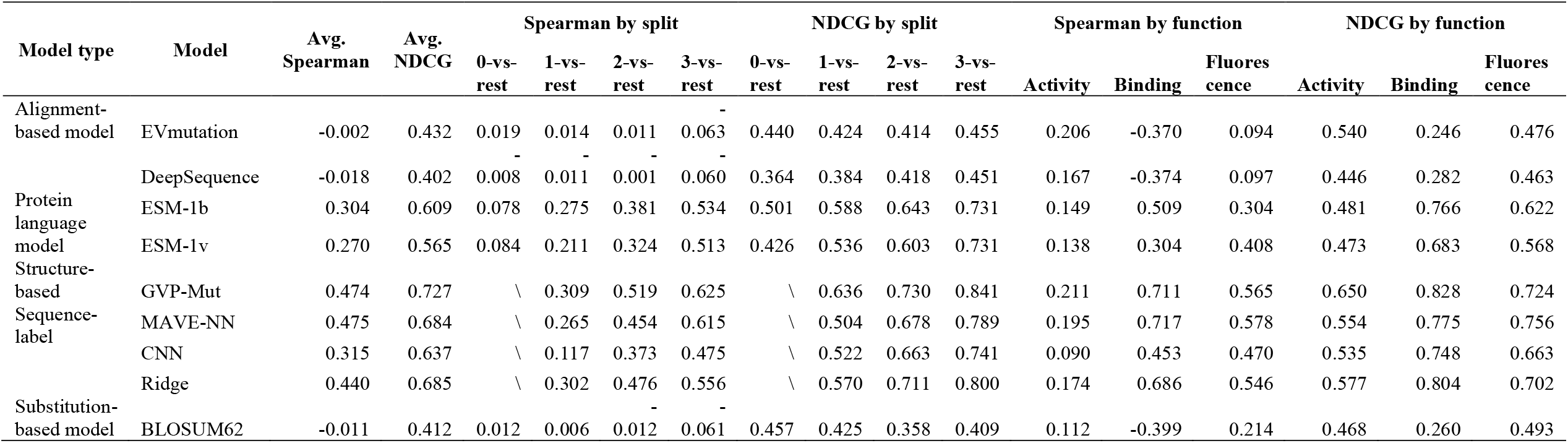
CombinGym benchmark summary.

**Figure 4.**
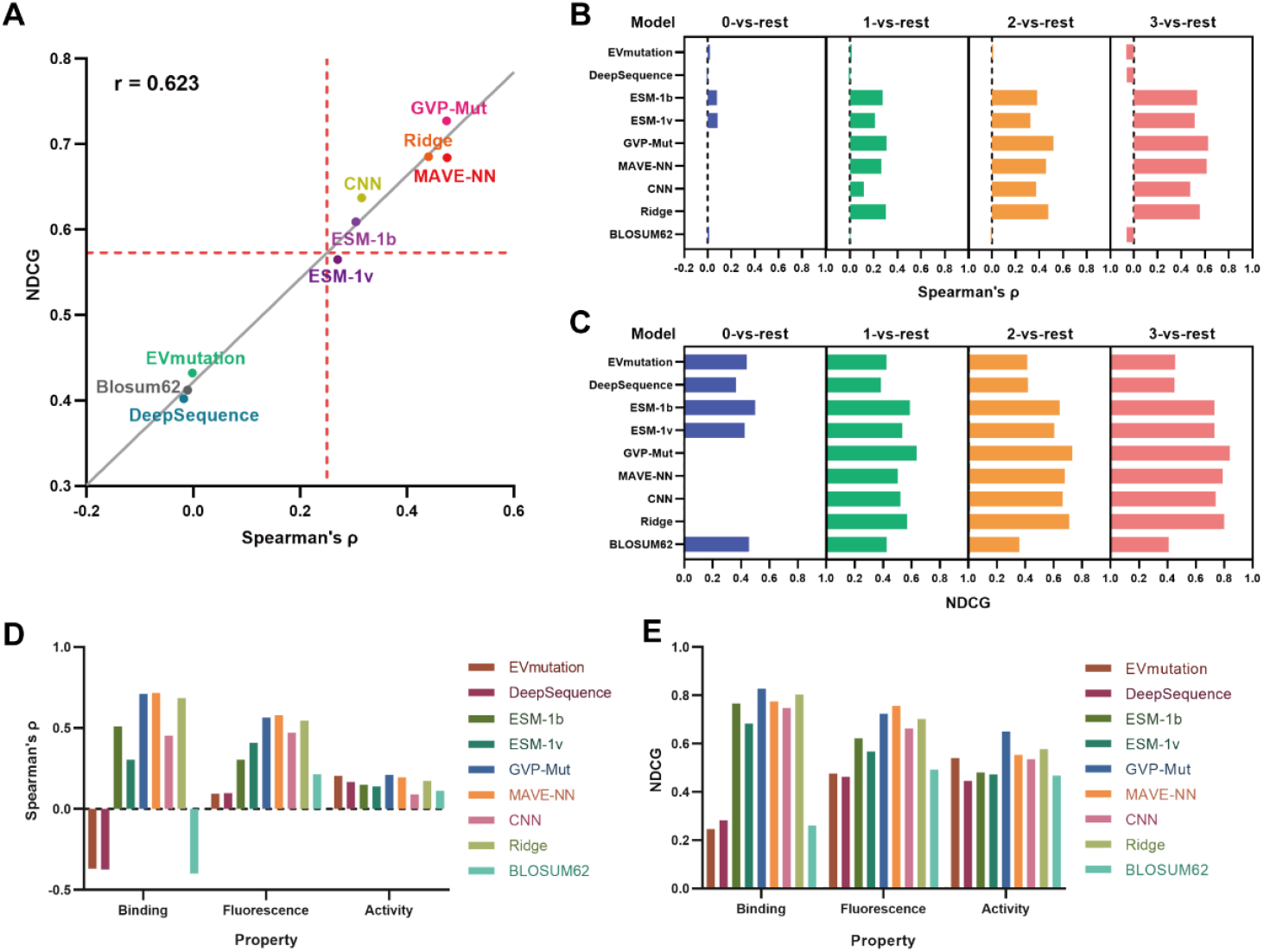
CombinGym benchmark. (**A**) Comparison of the overall performance of each model on Spearman’s ρ and NDCG. The solid gray line shows the linear regression of Spearman’s ρ and NDCG. The red dash lines indicate the mean value of Spearman’s ρ and NDCG of all models, respectively. (**B**-**C**) Comparison of the performance of each model in the setting of 0-vs-rest, 1-vs-rest, 2-vs-rest and 3-vs-rest for Spearman’s ρ (**B**) and NDCG (**C**). (**D**-**E**) Comparison of the performance of each model on the property of protein binding, fluorescence and enzymatic activity for Spearman’s ρ (**D**) and NDCG (**E**).

Among zero-shot prediction models, ESM-1b achieved slightly better predictive performance, though performance varied depending on specific protein or phenotypic traits. ESM-1b performed poorly on CreiLOV, the red fluorescence phenotype of eqFP611, and the total activity phenotype of RhlA, while EVmutation and DeepSequence showed superior performance in predicting CreiLOV fluorescence and RhlA activity. As the order of functional labels in the training set increased, supervised model performance improved, while unsupervised model performance deteriorated (Figure 4B and C). This observation indicates that predicting higher-order combinatorial mutants is more challenging than lower-order ones, but incorporating lower-order mutant data into the training set improves model performance. GVP-Mut and MAVE-NN showed the largest performance improvements and performed best on the 3-vs-rest task. Supervised models performed better in predicting protein binding and fluorescence properties than enzymatic activity (Figure 4D and E), indicating that enzymatic activity presents greater prediction challenges.

Multiple landscapes exhibiting distinct characteristics were included for bnAbs, eqFP611 and RhlA in CombinGym (Supplementary Figure S2). The fitness distributions for CR6261_H1 and CR6261_H9 were similar, and consistently, the Spearman’s ρ and NDCG metrics were comparable for most baseline models. The eqFP611 dataset contains three different but related landscapes: BFP, RFP and combined fluorescence. EVmutation and DeepSequence performed better for RFP in the 0-vs-rest setting, while ESM-1b and BLOSUM62 performed better for BFP and combined fluorescence. In supervised learning settings, Spearman’s ρ values followed the order BFP>RFP>combined fluorescence, while NDCG values for BFP and combined fluorescence were similar and higher than for RFP (Supplementary Figure S8). For RhlA, unsupervised models EVmutation and DeepSequence achieved better performance on total and specific activities, while ESM-1b and ESM-1v both exhibited poor performance across all three properties (Supplementary Figure S9). The strong performance of MSA-based algorithms on RhlA activity likely reflects evolutionary selection of functional sequences. However, in supervised settings, NDCG metrics were higher for specificity, indicating different properties of the same protein present distinct prediction challenges.

Spearman’s ρ and NDCG are common benchmarking metrics for protein prediction and design, respectively (28,55). Spearman’s ρ focuses on overall ranking consistency, while NDCG emphasizes accuracy for the top percent of variants. These metrics are positively correlated but not always. The two metrics in CombinGym show moderate correlation with a Pearson’s correlation coefficient of 0.623 (Figure 4A). We calculated the correlation between Spearman’s ρ and NDCG across different dimensions including protein property, model type and task scenarios. The correlation was lower for enzymatic activity than for protein binding and fluorescence (Supplementary Table S2). Among model types, the structure-based GVP-Mut model and sequence-label models exhibited relatively lower correlation coefficients.

### In silico engineering of CreiLOV higher-order mutants

We further explored whether CombinGym benchmark results could guide the extrapolative design of higher-order mutants using lower-order mutant data. We performed a simulation using the CreiLOV dataset which most models performed well. The objective was to design higher-order mutants with 4 to 15 mutations based on tested single, double, and triple mutants, so only supervised models were selected for training. We trained ESM-1b, ESM-1v, GVP-Mut, MAVE-NN, CNN, and Ridge models on the single and double mutants, predicted the fitness of all tested triple mutants, and calculated Spearman’s ρ and NDCG metrics. This approach is analogous to the 2-vs-rest scenario described above, enabling us to select the best models for the 3-vs-rest task. GVP-Mut, MAVE-NN, CNN and Ridge models reached better and similar performance on both Spearman’s ρ and NDCG metrics (Figure 5A and B). However, GVP-Mut required 24 times longer than CNN, 72 times longer than Ridge, and 360 times longer than MAVE-NN (Supplementary Table S3). ESM-1b, ESM-1v, and CNN had higher peak memory usage than GVP-Mut, followed by Ridge and MAVE-NN. Based on these considerations, MAVE-NN, CNN and Ridge were selected for predicting CreiLOV higher-order mutants.

**Figure 5.**
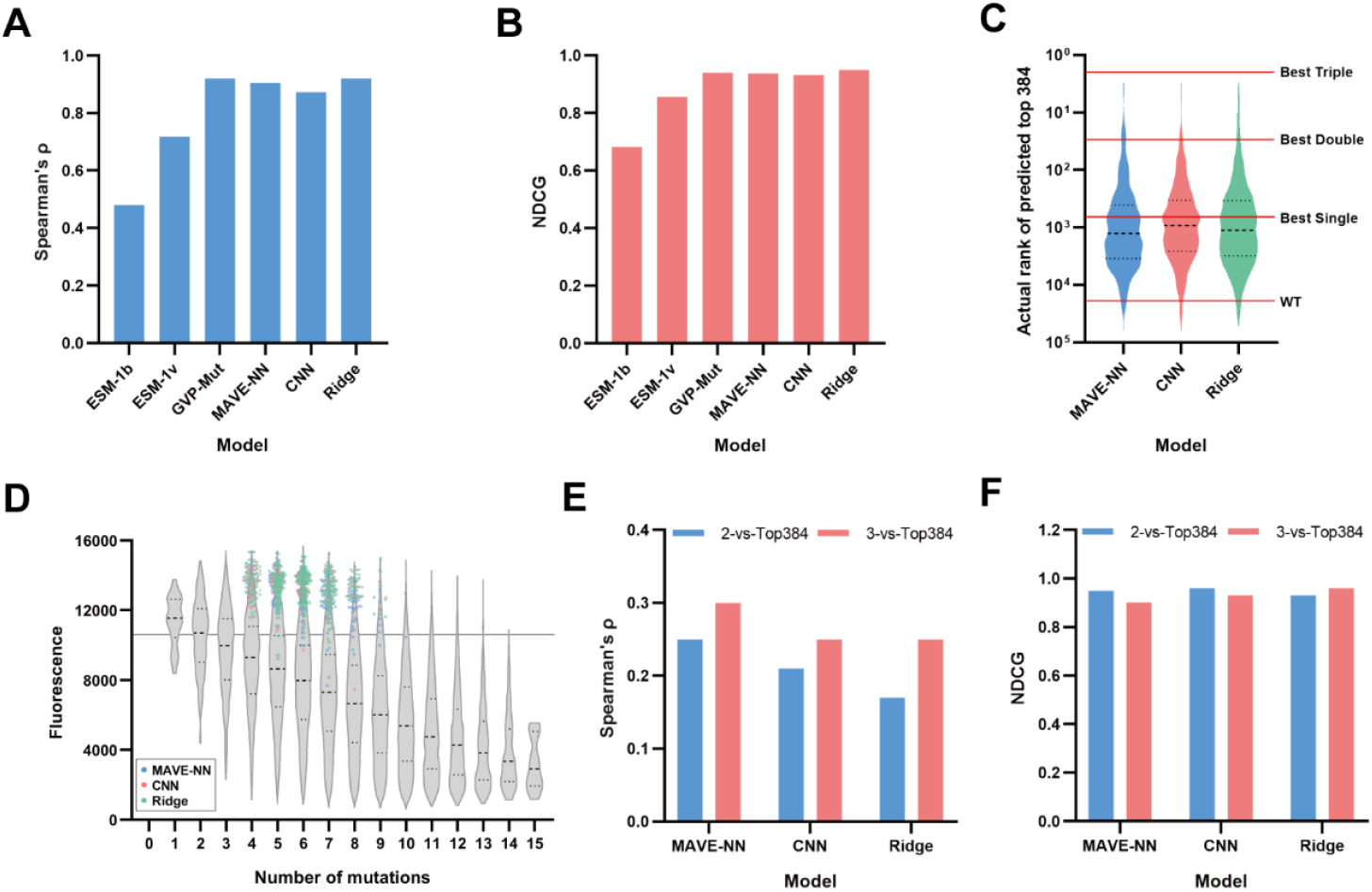
*In silico* engineering of CreiLOV higher-order mutants. (**A**-**B**) The Spearman’s ρ (**A**) and NDCG (**B**) of supervised models in the setting of 2-vs-3 for CreiLOV. (**C**) The actual rank of the top 384 CreiLOV mutants with more than three mutations predicted by indicated models using data of mutants with three mutations or less. The red solid lines show the rank of WT and best single, double and triple mutants. (**D**) The fluorescence distribution of predicted top 384 mutants overlapping with the overall fluorescence distribution of combinatorial library. The dash line indicates the fluorescence of WT CreiLOV. (**E**-**F**) Comparison of the Spearman’s ρ (**E**) and NDCG (**F**) for 2-vs-Top384 and 3-vs-Top384.

We prioritized the top 384 predictions per model for downstream analysis. Intersection analysis revealed that 157 mutants (40.9%) were common to all three models, and CNN and Ridge shared 263 overlapping predictions (68.5%). The predicted rankings spanned positions 3 to 61,426/64,394/52,675 across the 164,253 mutants, with mean ranks of 2,926/2,699/2,881 for MAVE-NN/CNN/Ridge, respectively. Approximately 98% of the predicted top 384 mutants were brighter than wild-type CreiLOV, 33%/38%/37% brighter than the best single mutant and 2.9%/2.1%/3.1% brighter than the best double mutant, respectively (Figure 5C). These mutants comprised of 4-9 amino acid substitutions with a modal number of 6 and were clustered within the top region of the fluorescence distributions (Figure 5D). When we compared performance of MAVE-NN, CNN, and Ridge models in the 2-vs-top384 and 3-vs-top384 settings, MAVE-NN outperformed CNN and Ridge on Spearman’s ρ, and all three models achieved better performance in the 3-vs-top384 scenario (Figure 5E). The NDCG values were similar for all models in both settings (Figure 5F), likely because model performance was already near optimal. The newly tested top 384 mutants likely provide additional information for machine learning, further improving model performance.

### Experimental engineering of RhlA higher-order mutants

To validate the practical utility of CombinGym, we conducted experimental protein engineering in a companion study (unpublished). Considering the overall ranking, NDCG values in the 2-vs-rest setting for RhlA, and computational requirements, we selected MAVE-NN for benchmarking. The phenotypes were predicted for all untested combinatorial mutants with 4-6 mutations. For the top candidate sequences from a protein design competition, we successfully constructed and tested variants using automated robotic DNA assembly and high-throughput mass spectrometry as previously reported (57). Comparative benchmarking across 0-3 and 0-6 datasets demonstrated consistent performance improvements in NDCG metrics (Figure 6). A dramatic improvement for specificity and specific activity was observed when triple mutants were added into training set. Our lower-order mutant dataset supported collaborative model development efforts in the field, achieving a substantial product yield improvement over the wild type.

**Figure 6.**
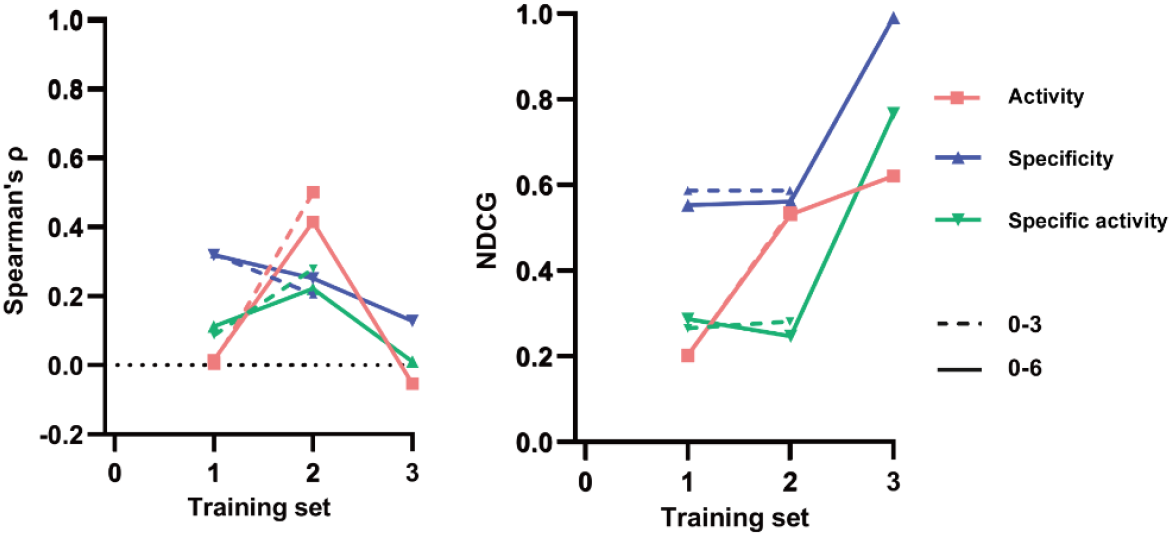
Experimental engineering of RhlA higher-order mutants. The comparison of MAVE-NN performance on Spearman’s ρ and NDCG for each phenotype using 0-3 or 0-6 dataset.

### CombinGym website

To promote dataset collaboration and model development, we developed the website of CombinGym (https://www.combingym.org), an online platform that hosts all datasets, benchmark scores, and model leaderboards. Each dataset entry contains a unique identification number (ID) and metadata (Table 3). Sequence information includes protein name, UniProt ID, wild-type sequence file, and sequence length. For DMS datasets, the mutation sites and their number, variant space, number and percentage of measured variants, DMS dataset file, and reference are included (Figure 7A). Currently, CombinGym contains 24 landscapes of 11 proteins, comprising 445 single, 5,027 double, 39,804 triple, and over 800,000 higher-order mutants. However, the datasets are heavily dominated by protein binding (76.65%), with enzymatic activity accounting for just 1.2% (Figure 7B). The RhlA enzyme dataset will be made available upon publication of the companion study. We also provide bitscore thresholds, MSA depth and sites, number of MSA sites and MSA files for each protein, along with AlphaFold3-predicted structures. Users can download all files through the website, as well as upload new experimental data with required metadata through a web form.

**Table 3.**
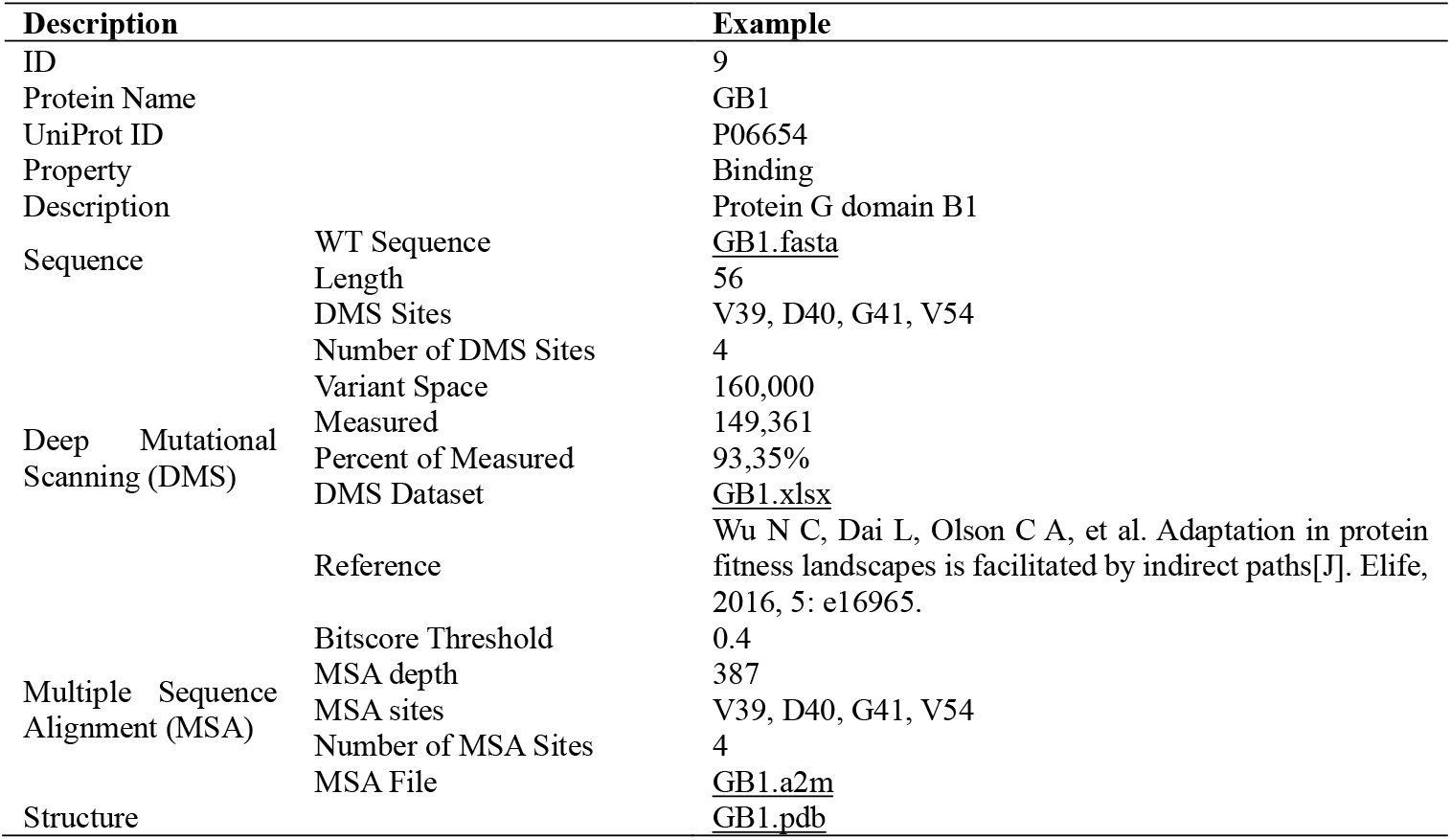
Description of data items in CombinGym dataset database with an example entry showing the GB1 dataset.

**Figure 7.**
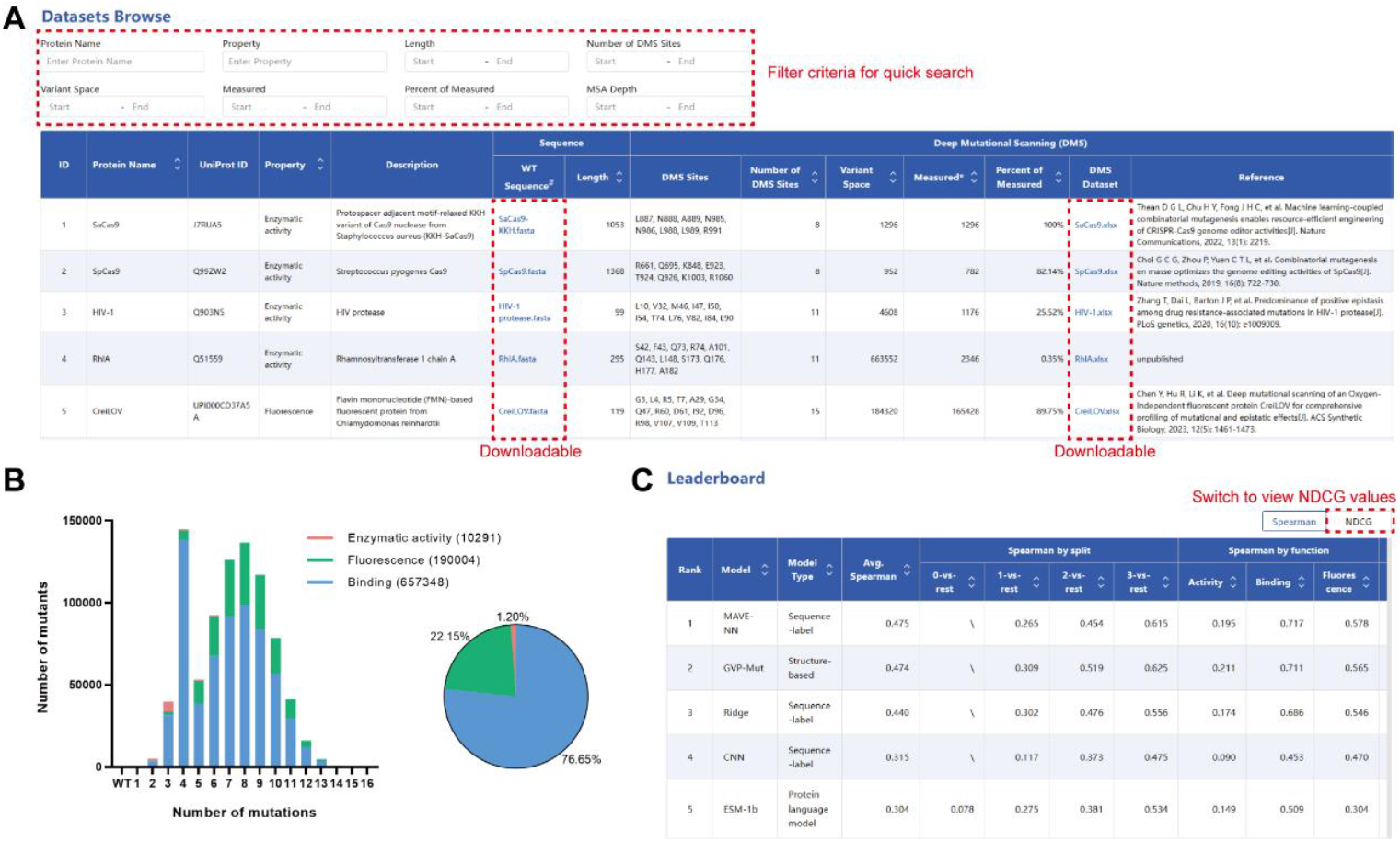
CombinGym combinatorial mutagenesis datasets, mutation statistics and model leaderboard. (**A**) Graphical interface of datasets for CombinGym. Above the data table shows filter criteria for quick search. The WT sequence and DMS dataset files are freely downloadable. (**B**) Mutation statistics of CombinGym. Left: Bar plot shows the number of mutants for each number of mutation and each type of protein property. Right: Pie plot shows the proportion of each protein property. (**C**) Graphical interface of model leaderboard for CombinGym. Switch to view NDCG values by clicking the ‘NDCG’ button.

The ‘Baselines’ page contains detailed benchmark scores for Spearman’s ρ and NDCG across each protein phenotype, baseline model, and scenario, totaling over 800 entries. A leaderboard ranks models for higher-order protein variant design, showing both overall performance and separate rankings for each split (0/1/2/3-vs-rest) and each functional category (enzyme activity, binding, and fluorescence) (Figure 7C). Through the ‘Service’ page, users can request automated protein engineering experiments via our integrated synthetic biology platform to validate model predictions or generate new experimental data. These services are supported by robust, high-throughput screening pipelines, as schematically illustrated in Supplementary Figure S10. Through community contribution mechanisms, CombinGym provides a unified platform for dataset sharing, model evaluation, and experimental validation, bridging the gap between computational predictions and experimental protein engineering.

## Discussion

In this study, we present CombinGym, a benchmarking platform for predicting and designing protein combinatorial mutants using machine learning models. To position CombinGym among existing benchmarks, we compared it with FLIP (27), ProteinGym (28), and FS-mutant (58), noting substantial differences in target tasks, dataset composition, experimental validation and data preprocessing approaches (Supplementary Table S4). First, to our knowledge, CombinGym is the first benchmark specifically designed for combinatorial mutagenesis, whereas existing benchmarks focus on single-residue or low-order mutants. Second, we analyzed how experimental noise and data preprocessing affect model performance, and these critical factors are rarely examined in other benchmarks. Third, it incorporates both *in silico* simulation and wet-lab validation, providing a practical testbed for real-world protein engineering applications. All datasets, benchmarks, and results are available through an interactive online database, facilitating community-driven expansion and transparent model comparison.

CombinGym currently benchmarks nine widely used models, but the field is rapidly evolving. Models explicitly designed for epistasis, such as Epistatic Net (67), ECNet (68) and MODIFY (69) merit future inclusion. It is also worth exploring whether hybrid architectures could better capture higher-order interactions by integrating multimodal data, including those combining alignment and protein language (TranceptEVE (59), PoET (60), Tranception (61), and MSA transformer (62)), as well as ProSST (63) that combines structure and protein language. Furthermore, Kermut (64) and ProteinNPT (65) are the top-performing supervised models while EVE (66) and ESM-2 (67) excel at zero-shot prediction for the ProteinGym benchmarking, which focuses on single-mutant prediction (28). Evaluating whether these models maintain performance for combinatorial variants would test generalization capabilities.

For protein engineers, CombinGym provides actionable guidance through a staged approach. First, target function strongly influences model selection. In zero-shot settings, MSA-based approaches yielded context-dependent performance: Spearman’s ρ values were negative for bnAbs and mTagBFP2 but positive for CreiLOV and RhlA. For supervised models, performance followed a clear hierarchy: protein binding > fluorescence > enzymatic activity, reflecting increasing biological complexity from direct molecular interactions to multi-step catalytic processes. This suggests researchers should pilot models on small datasets for their target function before committing to large-scale experiments, as demonstrated in our CreiLOV and RhlA validations (Figure 5 and 6). Second, third-order mutant data are critical for improving model prediction of higher-order variants (Figure 6), likely by capturing pairwise and three-way epistatic interactions that dominate protein fitness landscapes (12). Despite this importance, datasets containing triple mutants and beyond remain scarce (68-70) because the exponential increase in library size with additional sites poses practical limits on experimental scales. By interfacing with automated biofoundries (57,71-73), CombinGym provides a platform for community-driven collection and expansion of combinatorial variant data, as demonstrated by the RhlA case.

In conclusion, CombinGym establishes a comprehensive benchmarking framework for combinational protein mutagenesis, addressing critical gaps in current protein engineering research. By systematically evaluating model performance across diverse protein landscapes and providing both computational and experimental validation, CombinGym offers practical guidance for protein engineers which facilitating the development of next-generation machine learning models. The platform’s integration with automated biofoundries and community-driven expansion mechanisms positions it as a valuable resource for advancing the field of protein engineering through data-driven design and experimental validation.

