## Supplementary Information for "CombinGym: a benchmark platform for machine learning-assisted design of combinatorial protein variants"

#### Supplementary methods

##### Strains, media and cultivation conditions

*E. coli* DH5α cells were used for cloning and propagation of RhlA expression plasmids harboring wild type or combinatorial mutant sequences. *E. coli* BL21(DE3) cells were used for RLs production. Cells were cultured in Luria broth (LB) medium (10 g/L tryptone, 5 g/L yeast extract, 5 g/L NaCl, pH 7.0) supplemented with 50 μg/mL kanamycin at 37 °C and 220 rpm, or on 1.5% LB agar plates and incubated at 37 °C. For the inducible production of RLs, *E. coli* BL21(DE3) cells were cultured at 30 °C and supplemented with IPTG at a final concentration of 1 mM.

##### Construction of RhlA combinatorial mutant library

The RhlA combinatorial mutant library was constructed based on the insert plasmids prepared in our previous study. Briefly, we modularized the *rhlA* gene into four fragments and created inserts, respectively. Each of the inserts was incorporated into a level 0 receiver vector via *Bsm*BI-mediated Golden Gate assembly and was normalized to a final concentration of 30 ng/μL in a 384-well storage plate. To generate combinatorial mutant plasmids, the corresponding four insertions were selected with the Echo liquid-handling robot using a custom cherry-picking script and mixed with the level 1 receiver vector of pRSFDuet-*rhlb*-*ccdb* in 96-well PCR plates, followed by a Golden Gate assembly reaction using *Bsa*I. The mutation count in combinatorial mutants can be easily expanded by picking out the insertion plasmids with more mutations. The resulting reaction mixtures were chemically transformed into *E. coli* DH5α and 10~30 independent colonies were obtained in one well of 12-well plates. Sanger sequencing results indicated that a maximum of two clones is sufficient for correct construct.

##### RapidFire high-throughput mass spectrometry assay

For RL measurement, 500 µL of *E. coli* cultures were mixed with an equal volume of methanol, followed by centrifugation at 4,000 g for 10 minutes. The top phase was filtered through a 0.22 µm filter membrane and collected into a 96-well plate. A C18 cartridge (Agilent, #G9205A) was used for the RapidFire high-throughput MS system. Eluent A (acetonitrile:$H_2O$ (1:9, v/v)) and eluent B (acetonitrile:$H_2O$ (9:1, v/v)) mixed with 5 mM ammonium acetate were used as mobile phases. The mass spectrometer was equipped with an electrospray ionization (ESI) source and operated in negative mode with a capillary voltage of 5 kV and a cone voltage of 30 V. Nitrogen was used as the nebulizer gas, and the source temperature was maintained at 120 °C. Quantification was performed using the multiple reaction monitoring (MRM) mode.

#### Supplementary Figures

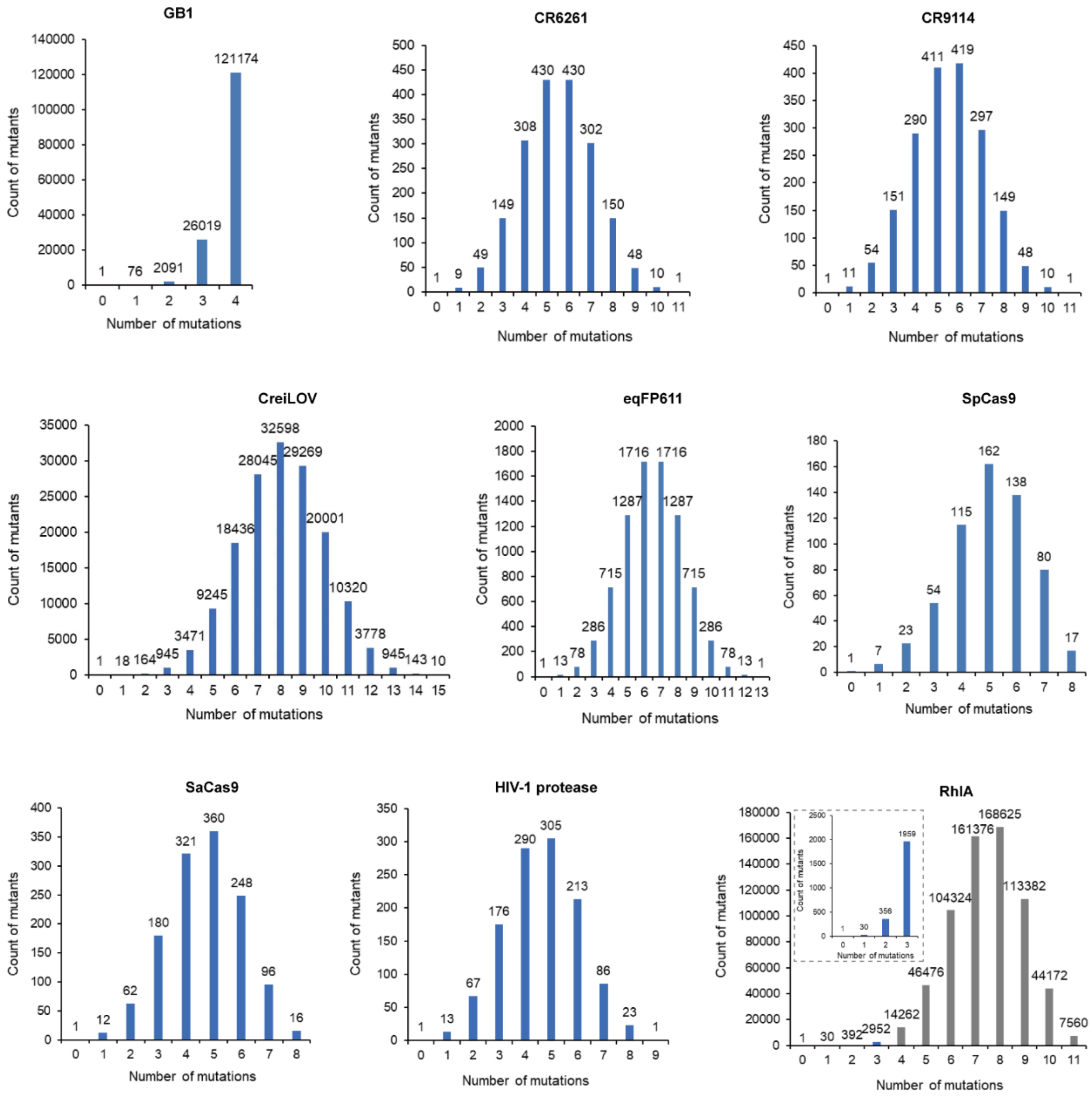


**Supplementary Figure S1**. The count distribution of the mutants with each number of mutations for all proteins. For RhlA, the gray bars show the untested higher-order mutants include in the local protein sequence space.

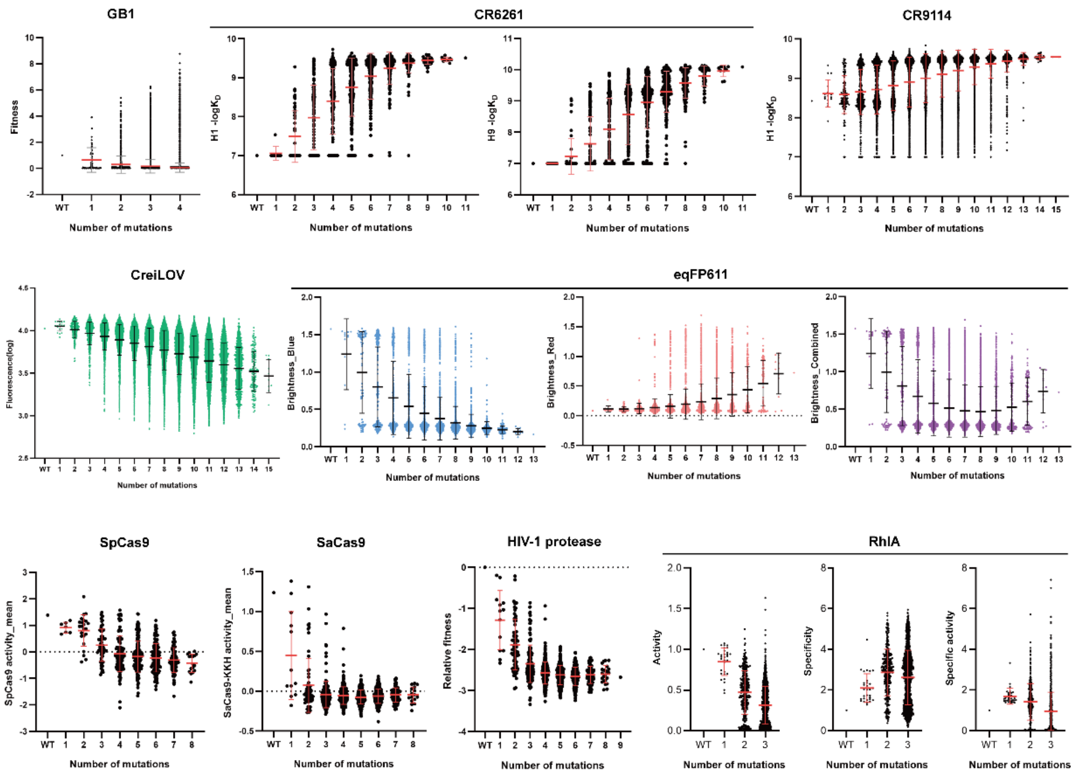


**Supplementary Figure S2**. The fitness distribution of the mutants with each number of mutations for all proteins. For CR6261, eqFP611 and RhlA, multiple landscapes are included.

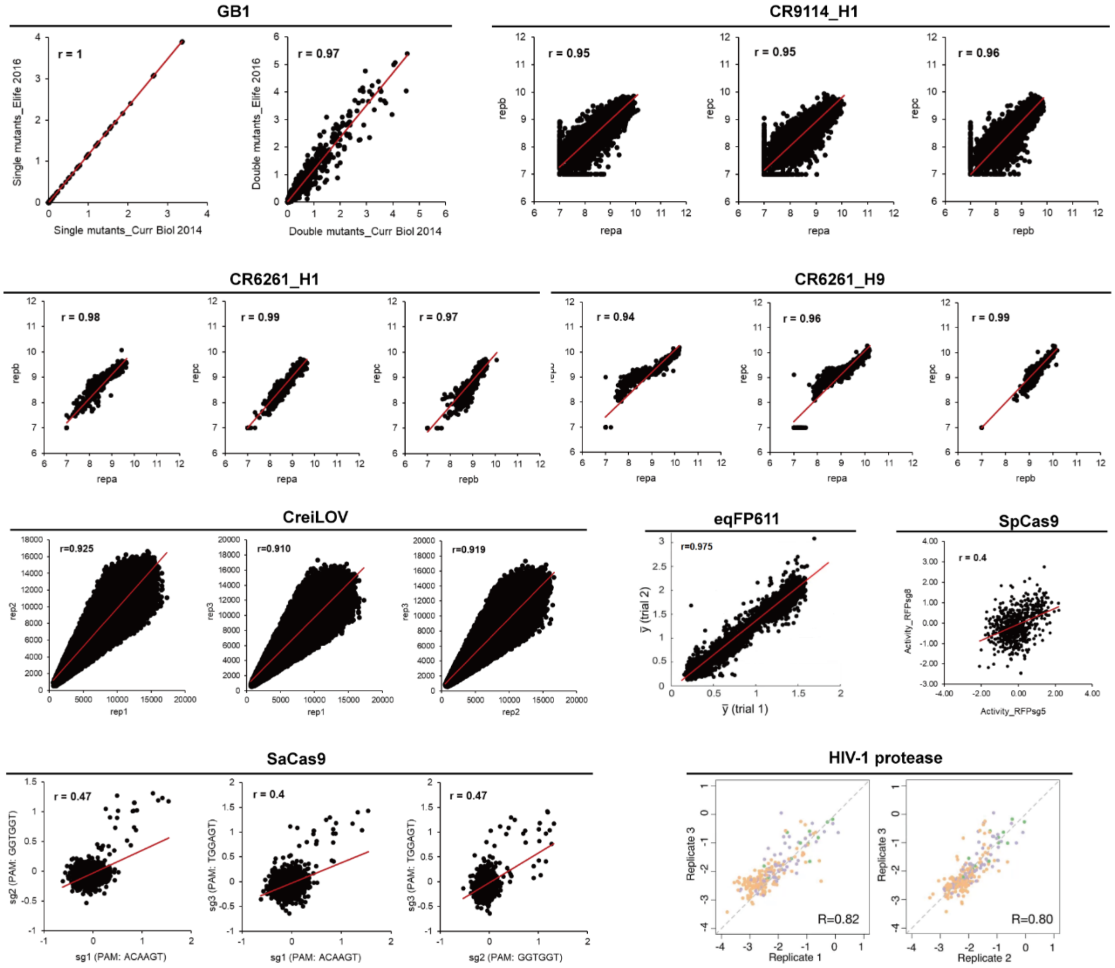


**Supplementary Figure S3**. The scatter plots and Pearson's correlation coefficients of the biological replicates for all landscapes.

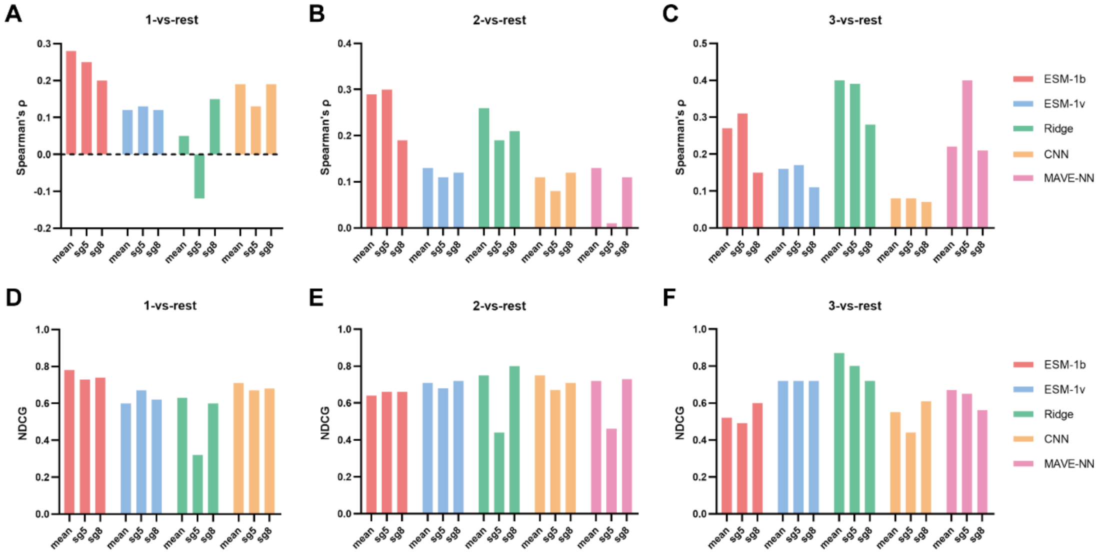


**Supplementary Figure S4**. The comparison of the benchmark of the supervised models trained by sg5, sg8 and the mean dataset of SpCas9. (**A-C**) The Spearman's ρ of each model in the setting of 1-vs-rest (**A**), 2-vs-rest (**B**) and 3-vs-rest (**C**), respectively. (**D-F**) The NDCG of each model in the setting of 1-vs-rest (**D**), 2-vs-rest (**E**) and 3-vs-rest (**F**), respectively.

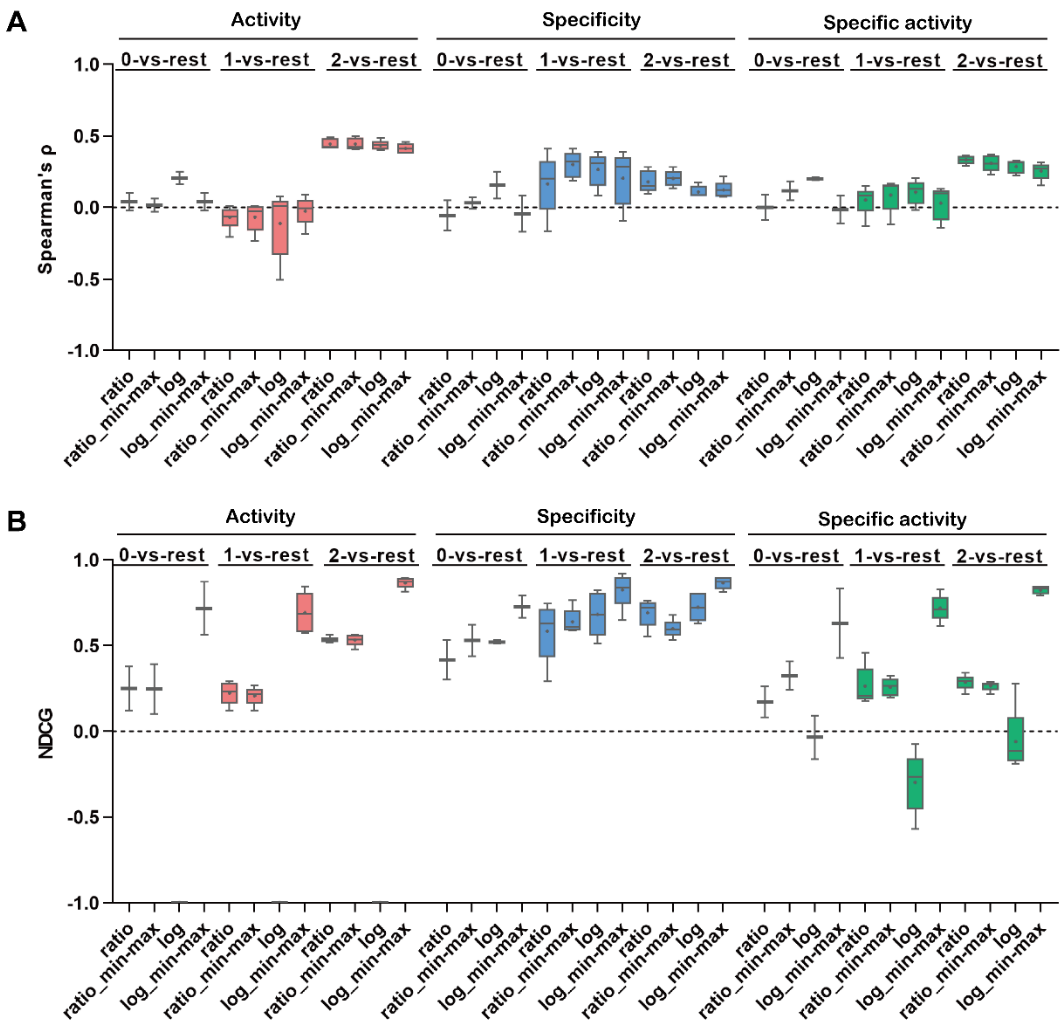


**Supplementary Figure S5**. The comparison of the overall benchmark of the models trained by the original ratio, min-max normalization, log normalization, or both log and min-max normalization datasets of RhlA. The Spearman's ρ (**A**) and NDCG (**B**) for activity, specificity and specific activity were obtained in the settings of 0-vs-rest, 1-vs-rest or 2-vs-rest, respectively.

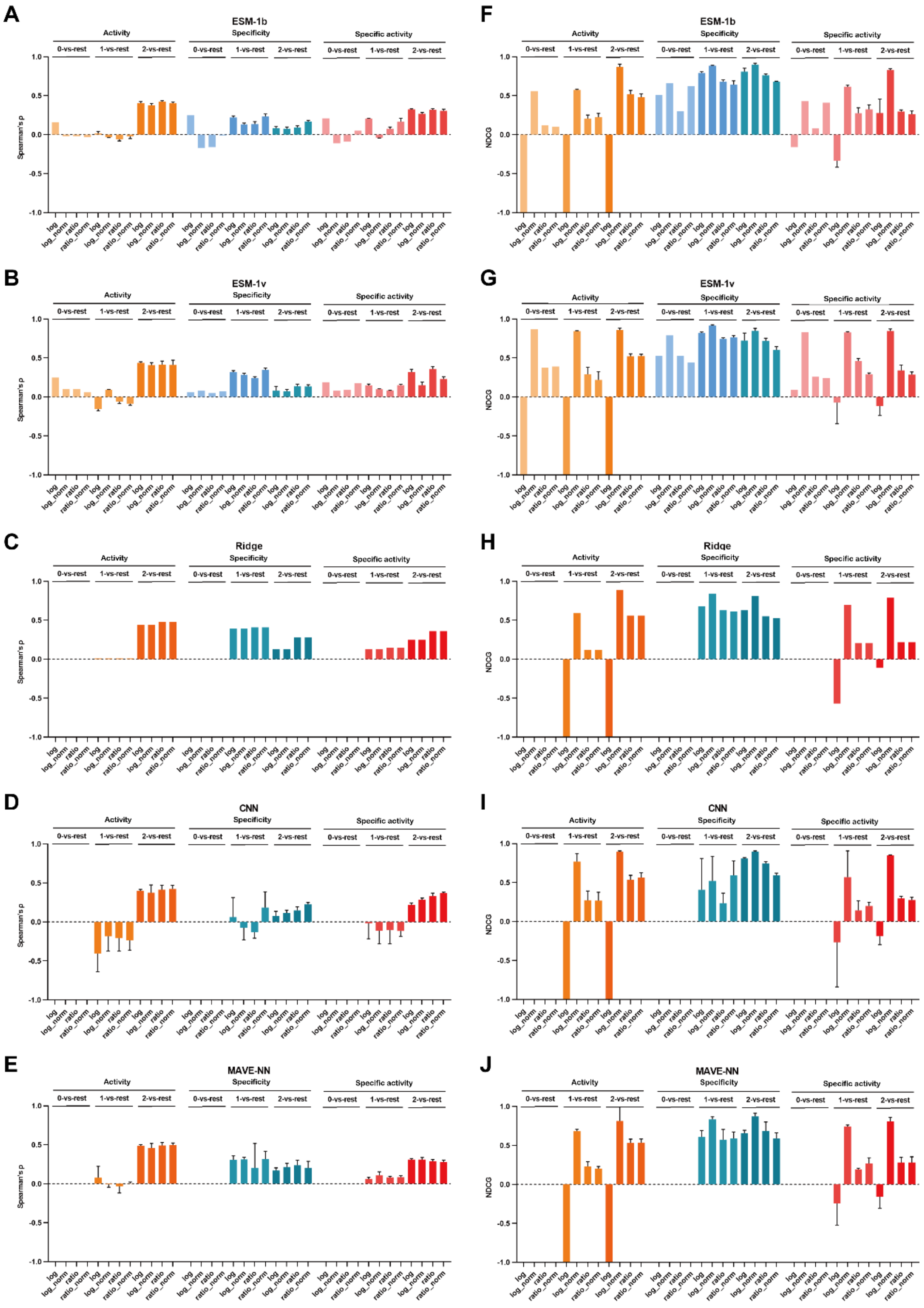


**Supplementary Figure S6**. The comparison of the detailed benchmark of the models trained by the original ratio, min-max normalization, log normalization, or both log and min-max normalization datasets of RhlA. The Spearman's ρ of ESM-1b (**A**), ESM-1v (**B**), Ridge (**C**), CNN (**D**) and MAVE-NN (**E**) and the NDCG of ESM-1b (**F**), ESM-1v (**G**), Ridge (**H**), CNN (**I**) and MAVE-NN (**J**) for activity, specificity and specific activity were obtained in the settings of 0-vs-rest, 1-vs-rest or 2-vs-rest, respectively.

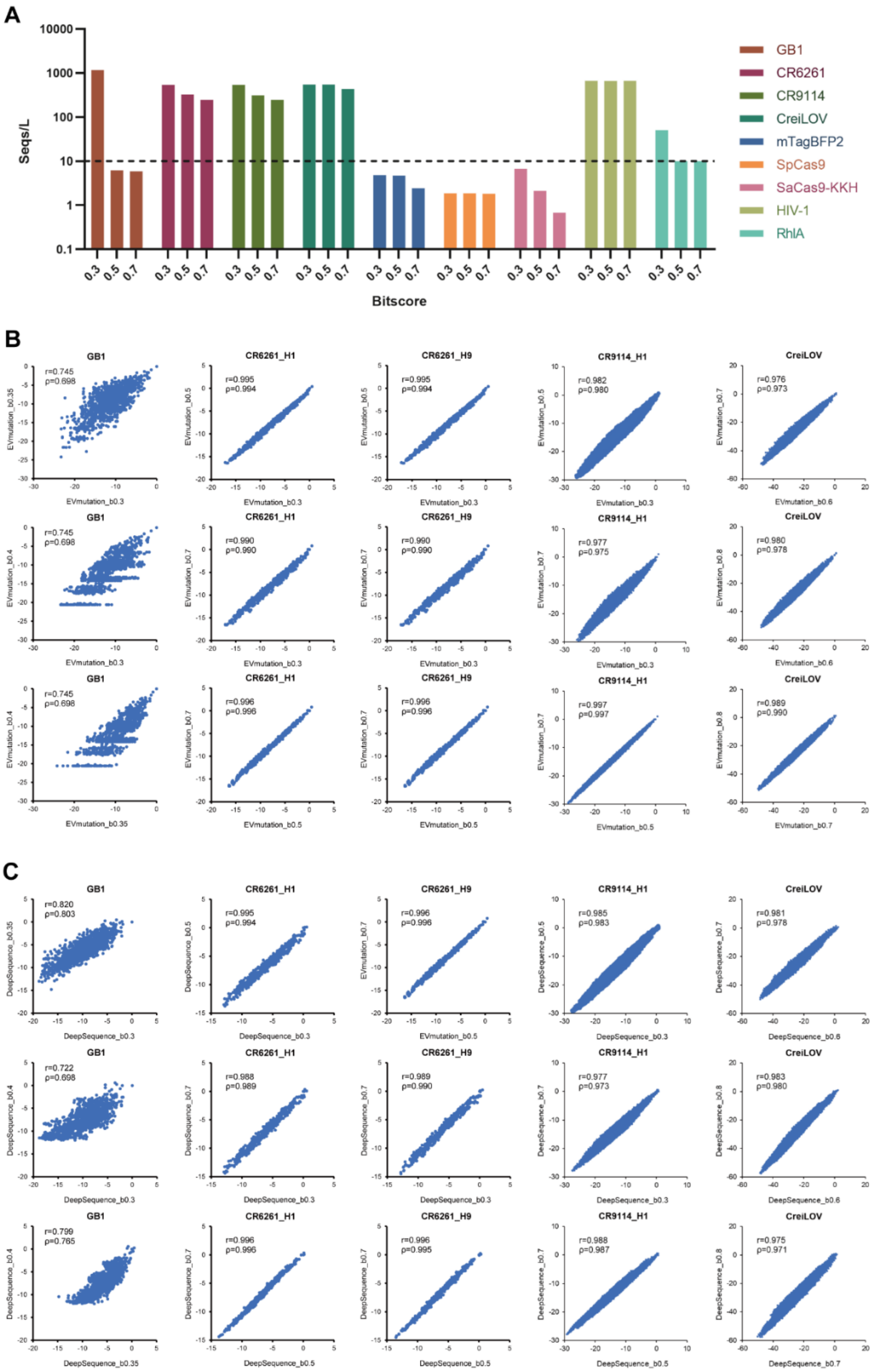


**Figure S7**. The impact of MSA-depth on the performance of alignment-based EVmutation and DeepSequence models. (**A**) The MSA depth for each protein obtained by default bitscore thresholds of 0.3, 0.5 and 0.7 using EVcouplings. (**B-C**) The correlation relationship of the fitness among different bitscore thresholds predicted by EVmutation (**B**) and DeepSequence (**C**) for selected GB1, CR6261, CR9114 and CreiLOV proteins.

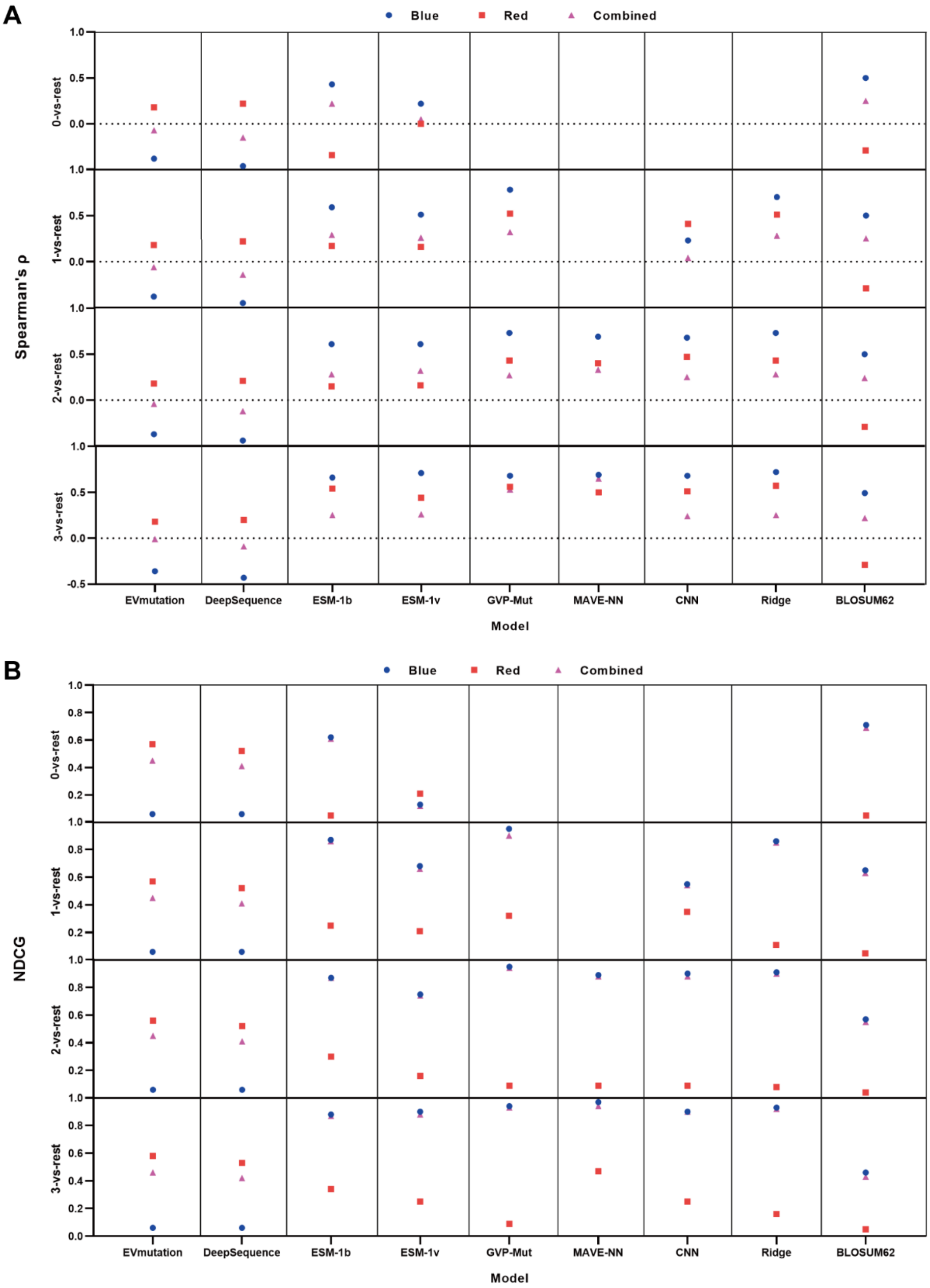


**Figure S8**. The comparison of the Spearman's ρ (**A**) and NDCG (**B**) metric values of the Blue, Red and Combined fluorescence phenotypes of eqFP611.

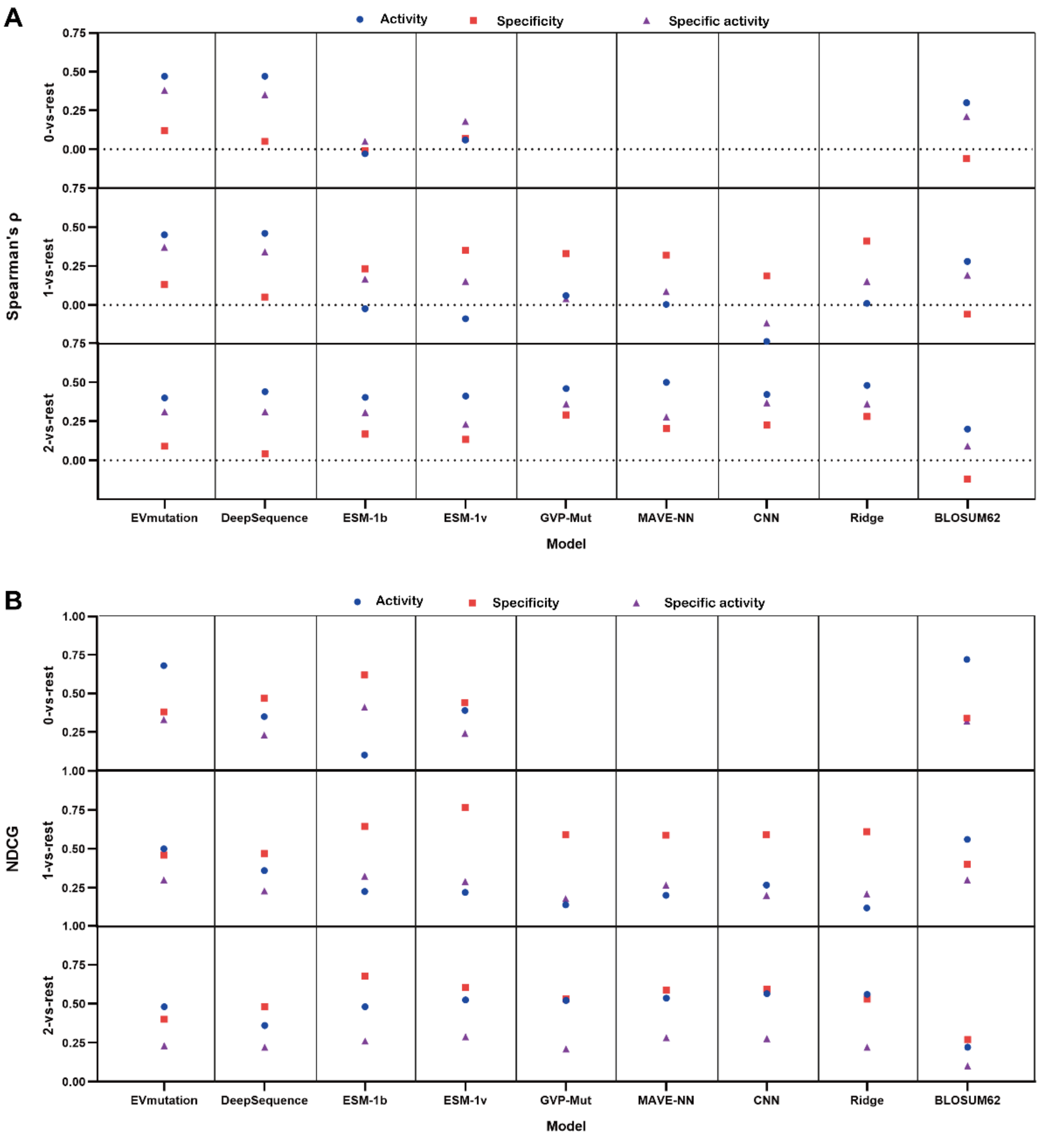


**Figure S9**. The comparison of the Spearman's ρ (**A**) and NDCG (**B**) metric values of activity, specificity and specific activity of RhlA.

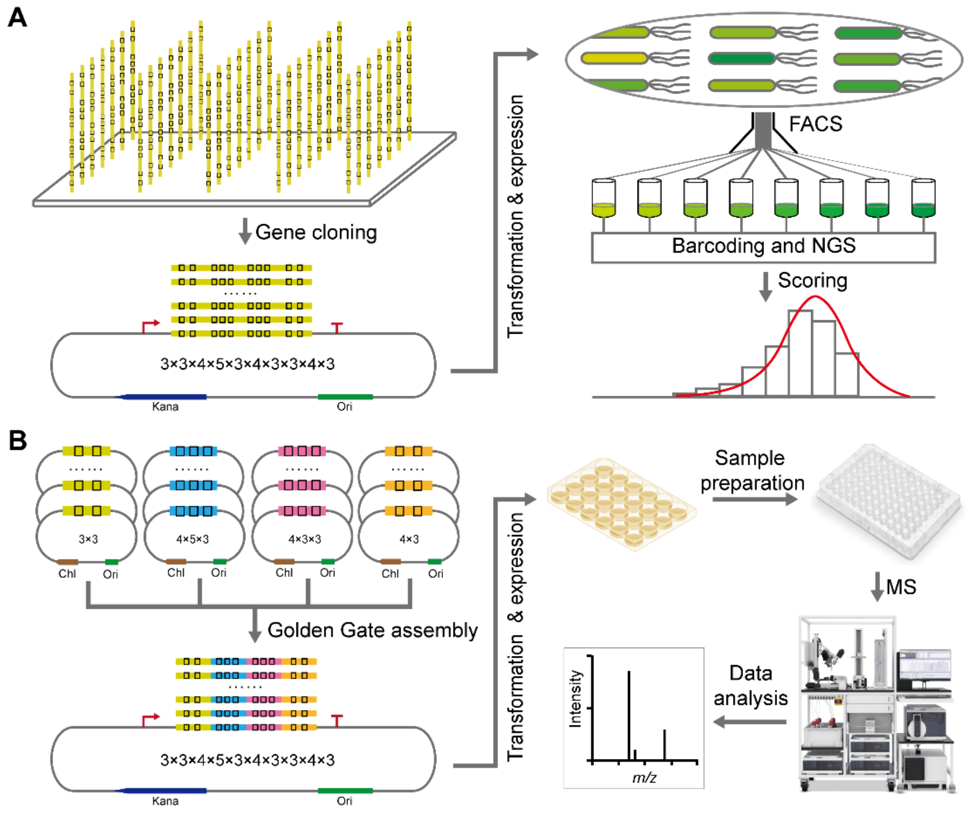


**Figure S10.** Scheme of high-throughput functional screening workflows for combinatorial protein libraries. (**A**) FACS-based screening workflow. The process begins with *in silico* designed DNA synthesis for library construction, followed by fluorescence-activated cell sorting (FACS) for screening, and culminates in high-throughput sequencing to link genotype to phenotype. (**B**) RapidFire mass spectrometry-based screening workflow. Library construction is achieved via automated and modular Golden Gate assembly, and functional screening is performed using high-throughput RapidFire mass spectrometry.

#### Supplementary Tables

**Supplementary Table S1**. Statistics of multiple sequence alignment by EVcouplings.

| query | bitscore_ threshold | minimum_column_ coverage | num_seqs | seq_len | seqs/L | num_cov | num_lc | perc_cov | 1st_uc | last_uc | len_cov | num_lc_ cov | N_eff |
|---|---|---|---|---|---|---|---|---|---|---|---|---|---|
| GB1 | 0.3 | 0.7 | 65489 | 56 | 1169.45 | 37 | 19 | 66.10% | 12 | 48 | 37 | 0 | 1455.117 |
| | 0.35 | 0.7 | 10511 | 56 | 187.70 | 33 | 23 | 58.90% | 18 | 50 | 33 | 0 | 768.265 |
| | 0.4 | 0.7 | 387 | 56 | 6.91 | 55 | 1 | 98.20% | 2 | 56 | 55 | 0 | 51.586 |
| | 0.45 | 0.7 | 346 | 56 | 6.18 | 55 | 1 | 98.20% | 2 | 56 | 55 | 0 | 28.672 |
| | 0.5 | 0.7 | 344 | 56 | 6.14 | 55 | 1 | 98.20% | 2 | 56 | 55 | 0 | 26.672 |
| | 0.7 | 0.7 | 328 | 56 | 5.86 | 55 | 1 | 98.20% | 2 | 56 | 55 | 0 | 23.422 |
| bnAbs_CR6261 | 0.3 | 0.7 | 65267 | 121 | 539.40 | 89 | 32 | 73.60% | 4 | 100 | 97 | 8 | 7930.157 |
| | 0.5 | 0.7 | 39017 | 121 | 322.45 | 97 | 24 | 80.20% | 3 | 100 | 98 | 1 | 4938.477 |
| | 0.7 | 0.7 | 29714 | 121 | 245.57 | 97 | 24 | 80.20% | 3 | 100 | 98 | 1 | 3914.841 |
| bnAbs_CR9114 | 0.3 | 0.7 | 65271 | 121 | 539.43 | 89 | 32 | 73.60% | 4 | 100 | 97 | 8 | 8001.269 |
| | 0.5 | 0.7 | 37683 | 121 | 311.43 | 97 | 24 | 80.20% | 3 | 100 | 98 | 1 | 4880.083 |
| | 0.7 | 0.7 | 29700 | 121 | 245.45 | 97 | 24 | 80.20% | 3 | 100 | 98 | 1 | 3912.666 |
| CreiLOV | 0.3 | 0.7 | 65512 | 119 | 550.52 | 99 | 20 | 83.20% | 16 | 114 | 99 | 0 | 36610.99 |
| | 0.4 | 0.7 | 65476 | 119 | 550.22 | 99 | 20 | 83.20% | 11 | 113 | 103 | 4 | 26222.42 |
| | 0.5 | 0.7 | 65493 | 119 | 550.36 | 106 | 13 | 89.10% | 7 | 113 | 107 | 1 | 16397.59 |
| | 0.6 | 0.7 | 65512 | 119 | 550.52 | 105 | 14 | 88.20% | 7 | 112 | 106 | 1 | 15006.45 |
| | 0.7 | 0.7 | 51241 | 119 | 430.60 | 106 | 13 | 89.10% | 7 | 112 | 106 | 0 | 10181.73 |
| | 0.8 | 0.7 | 33066 | 119 | 277.87 | 106 | 13 | 89.10% | 7 | 112 | 106 | 0 | 6504.347 |
| mTagBFP2 | 0.3 | 0.7 | 1118 | 233 | 4.80 | 210 | 23 | 90.10% | 5 | 221 | 217 | 7 | 165.605 |
| | 0.5 | 0.7 | 1093 | 233 | 4.69 | 211 | 22 | 90.60% | 4 | 221 | 218 | 7 | 151.438 |
| | 0.7 | 0.7 | 564 | 233 | 2.42 | 215 | 18 | 92.30% | 4 | 221 | 218 | 3 | 90.509 |
| SpCas9 | 0.15 | 0.7 | 6012 | 1368 | 4.39 | 796 | 572 | 58.20% | 367 | 1242 | 876 | 80 | 1835.474 |
| | 0.2 | 0.7 | 2573 | 1368 | 1.88 | 1279 | 89 | 93.50% | 3 | 1362 | 1360 | 81 | 581.995 |
| | 0.25 | 0.7 | 2573 | 1368 | 1.88 | 1280 | 88 | 93.60% | 3 | 1362 | 1360 | 80 | 579.266 |
| | 0.3 | 0.7 | 2564 | 1368 | 1.87 | 1284 | 84 | 93.90% | 3 | 1362 | 1360 | 76 | 572.638 |
| | 0.5 | 0.7 | 2559 | 1368 | 1.87 | 1282 | 86 | 93.70% | 3 | 1362 | 1360 | 78 | 572.02 |
| | 0.7 | 0.7 | 2496 | 1368 | 1.82 | 1280 | 88 | 93.60% | 3 | 1362 | 1360 | 80 | 533.109 |
| SaCas9-KKH | 0.3 | 0.7 | 6990 | 1053 | 6.64 | 761 | 292 | 72.30% | 5 | 845 | 841 | 80 | 2281.766 |
| | 0.5 | 0.7 | 2236 | 1053 | 2.12 | 755 | 298 | 71.70% | 6 | 802 | 797 | 42 | 609.737 |
| | 0.7 | 0.7 | 711 | 1053 | 0.68 | 1031 | 22 | 97.90% | 4 | 1048 | 1045 | 14 | 180.979 |
| HIV-1 protease | 0.3 | 0.7 | 65535 | 99 | 661.97 | 99 | 0 | 100.00% | 1 | 99 | 99 | 0 | 22.336 |
| | 0.5 | 0.7 | 65534 | 99 | 661.96 | 99 | 0 | 100.00% | 1 | 99 | 99 | 0 | 3 |
| | 0.7 | 0.7 | 65535 | 99 | 661.97 | 99 | 0 | 100.00% | 1 | 99 | 99 | 0 | 3 |
| RhlA | 0.3 | 0.7 | 14830 | 295 | 50.27 | 246 | 49 | 83.40% | 21 | 272 | 252 | 6 | 4320.547 |
| | 0.5 | 0.7 | 2957 | 295 | 10.02 | 278 | 17 | 94.20% | 1 | 278 | 278 | 0 | 114.448 |
| | 0.7 | 0.7 | 2957 | 295 | 10.02 | 278 | 17 | 94.20% | 1 | 278 | 278 | 0 | 113.876 |

**Supplementary Table S2**. Pearson's correlation coefficient between Spearman's ρ and NDCG metrics.

| Protein property/Model type/Task | | Pearson's r |
|---|---|---|
| Overall | | 0.623 |
| Protein property | binding | 0.697 |
| | fluorescence | 0.717 |
| | activity | 0.217 |
| Model type | Alignment-based | 0.557 |
| | Protein language model | 0.631 |
| | Structure-based | 0.388 |
| | Sequence-label | 0.444 |
| | Substitution-based | 0.617 |
| Task | 0-vs-rest | 0.596 |
| | 1-vs-rest | 0.583 |
| | 2-vs-rest | 0.559 |
| | 3-vs-rest | 0.658 |

**Supplementary Table S3**. Comparisons of time cost and memory usage.

| Model | Time | Peak Memory Usage | Spearman's ρ | NDCG |
|---|---|---|---|---|
| ESM-1b | 8 min | 10 G | 0.48# | 0.68# |
| ESM-1v | 8 min | 10 G | 0.72# | 0.88# |
| GVP-Mut | 120 min* | 2.5 G | 0.92 | 0.94 |
| MAVE-NN | 0.33 min | 0.5 G | 0.91 | 0.94 |
| CNN | 5 min | 14 G | 0.87 | 0.93 |
| Ridge | 1.66 min | 0.5 G | 0.92 | 0.95 |

### Weaker performance

* Larger time cost

**Supplementary Table S4**. Comparison with existing benchmarks.

| Benchmark | Target Task(s) | Dataset Type | Data Volume | Data Preprocessing benchmark | Experimental Validation | Online Database | Reference |
|---|---|---|---|---|---|---|---|
| FLIP | Fitness prediction | DMS and proteome, single and multiple mutants | ~300K variants | mutational depth | No | No | Dallago C, Mou J, Johnston K E, et al. FLIP: Benchmark tasks in fitness landscape inference for proteins[J]. bioRxiv, 2021: 2021.11. 09.467890. |
| ProteinGym | Fitness prediction and protein design | DMS and clinical, mostly single mutants | >2.7M variants | MSA depth, mutational depth, taxa, assayed phenotype | No | Yes | Notin P, Kollasch A, Ritter D, et al. Proteingym: Large-scale benchmarks for protein fitness prediction and design[J]. Advances in Neural Information Processing Systems, 2024, 36 |
| FS-mutant | Few-shot prediction | DMS, single and multiple mutants | >1.2M variants | mutational depth, training set size | No | No | Li M, Yu H, Fan G, et al. FS-mutant: A Few-shot Learning Benchmark for Protein Mutants Mining[J]. 2024. |
| CombinGym | Higher-order combinatorial mutant prediction and design | DMS, comprehensive single and multiple mutants | >800K variants | MSA depth, mutational depth, assayed phenotype, normalization method, measurement noise | Yes | Yes | This study |
